# Ovarian cancer-on-chip for patient-specific profiling of treatment responses to sequential chemotherapy and PD-L1 blockade

**DOI:** 10.64898/2026.07.21.739796

**Authors:** Sarah Plöger, Nicole Anderle, Eileen Wegner, Tobias Schmidt, Julia Roosz, Tengku Ibrahim Maulana, Lena Christ, Tobias Engler, Andreas Hartkopf, André Koch, Sara Y. Brucker, Katja Schenke-Layland, André Rosa, Christian Schmees, Peter Loskill

## Abstract

**Background:** Ovarian cancer (OvCa) ranks as the most lethal gynecological malignancy in women worldwide. This complex disease, which can develop independently of a woman’s age, is characterized by late diagnosis, pronounced tumor heterogeneity, and an immunosuppressive tumor microenvironment (TME). Incremental diagnostic tools that could better inform clinicians on potential therapy resistance or subsets of patients that could benefit from new drug modalities represent a critical unmet need to improve patient care and potentially the identification of new biomarkers.

**Objective:** This study aimed to establish a reconfigurable patient-derived OvCa-on-chip platform for longitudinal functional profiling of tumor cell death, immune activation, and patient-specific responses to TIL-mediated killing, PD-L1 blockade, and sequential chemo-immunotherapy.

**Methods:** Patient-derived OvCa microtumors (PDM) were integrated with sequential integration of autologous tumor-infiltrating lymphocytes (TILs) into a perfusable microfluidic chip in the presence of different single and combination treatment regimens of chemotherapy and immune checkpoint inhibitors (ICIs). Treatment responses were assessed by longitudinal quantification of caspase-cleaved cytokeratin 18 (ccCK18) as marker of apoptotic epithelial tumor cell death, as well as cytokine/chemokine release in chip effluents, and multiplex flow cytometry-based characterization of autologous TIL subsets.

**Results:** The perfusable OvCa-on-chip platform supported long-term culture of PDM while maintaining key structural and microenvironmental features of the primary tumor. Multidimensional analyses, including time-resolved assessment of tumor cell death, secretome profiling and correlative analysis of autologous TIL subsets revealed patient-specific tumor-immune response patterns and heterogenous sensitivity to TIL-mediated killing, PD-L1 blockade, and sequential chemo-immunotherapy. Correlation analyses identified treatment-dependent associations between specific TIL phenotypes and functional tumor cell killing. PD-1-expressing CD4⁺ TIL subsets correlated with enhanced tumor cell killing, whereas terminally exhausted CD8⁺PD-1⁺Tcf1⁻ TILs negatively correlated with durvalumab responses. In contrast, tumor-reactive CD8⁺CD39⁺ TILs were associated with improved responses under sequential chemo-immunotherapy conditions.

**Conclusion:** Collectively, this OvCa-on-chip system represents a complex in vitro model (CIVM) that combines 3D tumor tissue with autologous immune cells in a microfluidic platform. Resembling a physiologically relevant human preclinical platform, it allows for the time-resolved functional assessment of patient-specific responsiveness to OvCa therapies, with direct implications for personalized treatment stratification.

## Background

Although most patients with ovarian cancer (OvCa) initially respond to standard of care treatment, comprising of debulking surgery combined with platinum- and taxane-based chemotherapy, it remains the most lethal gynecologic malignancy (1). High-grade serous ovarian carcinoma (HGSOC), accounting for 70-80% of epithelial OvCa, is the predominant subtype, associated with aggressive disease progression, high relapse rates, and poor overall survival (1). The majority of patients lack specific symptoms and are often diagnosed at advanced stages of the disease, when the cancer has already metastasized (1). Approximately 75% of advanced-stage cases relapse within three years and frequently develop chemoresistance leading to a five-year survival rate below 40% (1, 2). Mainly, the pronounced inter- and intratumoral heterogeneity is considered to contribute to therapy resistance, frequent relapse, and limited treatment options in recurrent OvCa (3). Given the clinical success of immune checkpoint inhibitors (ICI) in several other cancer types, ICI-based strategies have also been studied in OvCa. However, ovarian cancer generally exhibits limited benefit from immunotherapeutic approaches such as PD-1/PD-L1 blockade likely reflecting the highly variable and often immunosuppressive immune landscape of the OvCa tumor microenvironment (TME) marked by immunosuppressive activity, impaired antitumor immunity, lack of immune infiltration and insufficient effective T-cell responses (4). Recent spatial and single-cell profiling studies have demonstrated that HGSOC is characterized by pronounced intra- and intertumoral immune heterogeneity, with distinct tumor subclones and anatomical sites shaping local immune evasion mechanisms and therapy resistance (5). Chemotherapy treatment may further reshape the TME landscape by enhancing immune cell infiltration and T-cell activation to potentiate the immunogenicity or by enhancing expression of immune checkpoint pathways (6). This highlights the need to better understand tumor-immune interactions during treatment to enable reliable biomarker identification for patient stratification. Together, these features establish OvCa as a highly relevant model for investigating determinants of heterogeneous responses to immunotherapy approaches in a clinically relevant immune-desert and immune-excluded tumor type.

Critical aspects of tumor-immune cell interactions, immunosuppressive TME signaling, and the temporal evolution of therapy resistance are often underrepresented or entirely absent in current preclinical cancer models providing limited translational value (7). Mainly because standard two-dimensional (2D) cell culture systems and many animal models fail to adequately recapitulate human tumor biology, the native tissue architecture, TME landscape, and therapy-induced dynamic responses (7). Addressing this complexity requires *ex vivo* preclincal models that can capture the physiologically relevant three-dimensional (3D) tumor architecture with preserved native TME components and the patient-specific tumor-immune crosstalk under treatment-relevant, dynamic conditions (7, 8). To address this challenge, we previously established patient-derived microtumors (PDM) as a 3D preclinical *ex vivo* tumor model that can be isolated together with autologous tumor-infiltrating lymphocytes (TILs) within two days post-tissue collection from a variety of fresh patient tumor tissue, including OvCa (9-13). Importantly, OvCa PDM were shown to be viable in static culture, preserve key histopathological and molecular characteristics of the original tumor (9). In this study, we present a novel microfluidic chip system that integrates OvCa PDM with a patient-matched immune compartment. This versatile platform enables longitudinal monitoring of functional tumor-immune interactions and treatment responses within a patient-specific TME. Our approach follows the implementation of sequential treatment regimes, consisting of chemotherapy, with the perfusion of paclitaxel followed by the time-controlled administration of autologous TILs, alone or in combination with the PD-L1 inhibitor Durvalumab (DU). Using this immunocompetent platform, we were able to longitudinally assess tumor viability, treatment-induced and TIL-induced tumor cell death, synergistic treatment responses, functional immune activation states, as well as cytokine/chemokine secretion dynamics. Summarized, this microfluidic OvCa system provides a physiologically relevant framework for investigating patient-specific tumor-immune interactions, treatment-induced immune modulation, and heterogeneous responses to ICI-based therapeutic strategies.

## Methods

### Human Specimens

Ovarian tumor tissue was collected from six OvCa patients undergoing surgery at the Center for Women’s Health, University Hospital Tübingen. Written informed consent was obtained and the project was approved by the local ethics committee (703/2019BO2). Tumors were categorized according to International Federation of Gynecology and Obstetrics (FIGO) grading system. Available clinical follow up data is provided in Table S1.

### PDM and TIL isolation

OvCa PDM and autologous TILs, were isolated as described before (9, 10).

### Immunohistochemical staining

Hematoxylin and eosin (HCE) and 3,3’-diaminobenzidine (DAB) staining was performed on formalin-fixed, paraffin-embedded (FFPE) OvCa PDM sections. OvCa PDM were fixed in 4% Roti® Histofix (Carl Roth) for 1 h at RT, incubated in Harris hematoxylin (Leica Biosystems) for 5 min, washed in dH2O, and dehydrated in four ethanol incubation steps (2x 50% ethanol, 2x 70% ethanol, 15 min each). OvCa PDM were embedded in Richard-Allan Scientific™ HistoGel™ (Thermo Fisher Scientific) using Tissue-Tek® Cyomolds® (Sakura). Tissue processing was performed using the HistoCore PEARL (Leica Biosystems). Subsequently, OvCa PDM processed Histogel blocks were embedded in paraffin and three micrometer sections were cut. Immunohistochemical staining was performed using the Dako Autostainer Link 48 (Agilent). Dako PT Link (Agilent) was used for antigen retrieval according to the manufacturer’s protocol. Stained FFPE slides were imaged using the Axio Scan Z1. All primary antibodies were validated on custom tissue sections. A comprehensive overview of the applied antibodies can be found in Table S2.

### Microfluidic chip fabrication

The OvCa PDM chip consists of two Polydimethylsiloxane (PDMS) outer layers. The tissue layer is made up of four tissue chambers, each measuring 1mm x 1mm x 1mm, with the diameter of the chamber widening towards the top, and an inlet and outlet for the media perfusion. The media channel layer, which is designed to provide a constant flow of media beneath the tissue chambers and membrane (Fig. 1B). The two PDMS layers are separated by a porous Polyethylene terephthalate (PET) membrane (pore size 3 µm) forming a porous barrier between the media channel and the tissue chambers. Membrane activation was achieved by plasma-enhanced chemical vapor deposition (PECVD), as previously described in (14). To produce the layers, PDMS was homogeneously mixed 10+1 with curing agent, degassed, and poured into custom molds. The layers were baked at 60 °C for a minimum of 4 h for the tissue layer or a minimum of 2 h for the media channel layer. subsequently, PDMS layers were rinsed with isopropanol and dried with compressed air. The PDM chips were assembled in two consecutive bonding steps by exposition to oxygen plasma activation (75 W, 0.2 cm^3^m−1 O_2_; Diener Pico, Diener electronic GmbH + Co. KG, Ebhausen, Germany). Firstly, the membrane was bonded to the tissue layer and backed at 60 °C for 1 h to reverse the plasma activation. Secondly, the media channel layer was bonded to the tissue layer containing the PET membrane. The assembled chip was backed at 60 °C overnight.

**Fig. 1:**
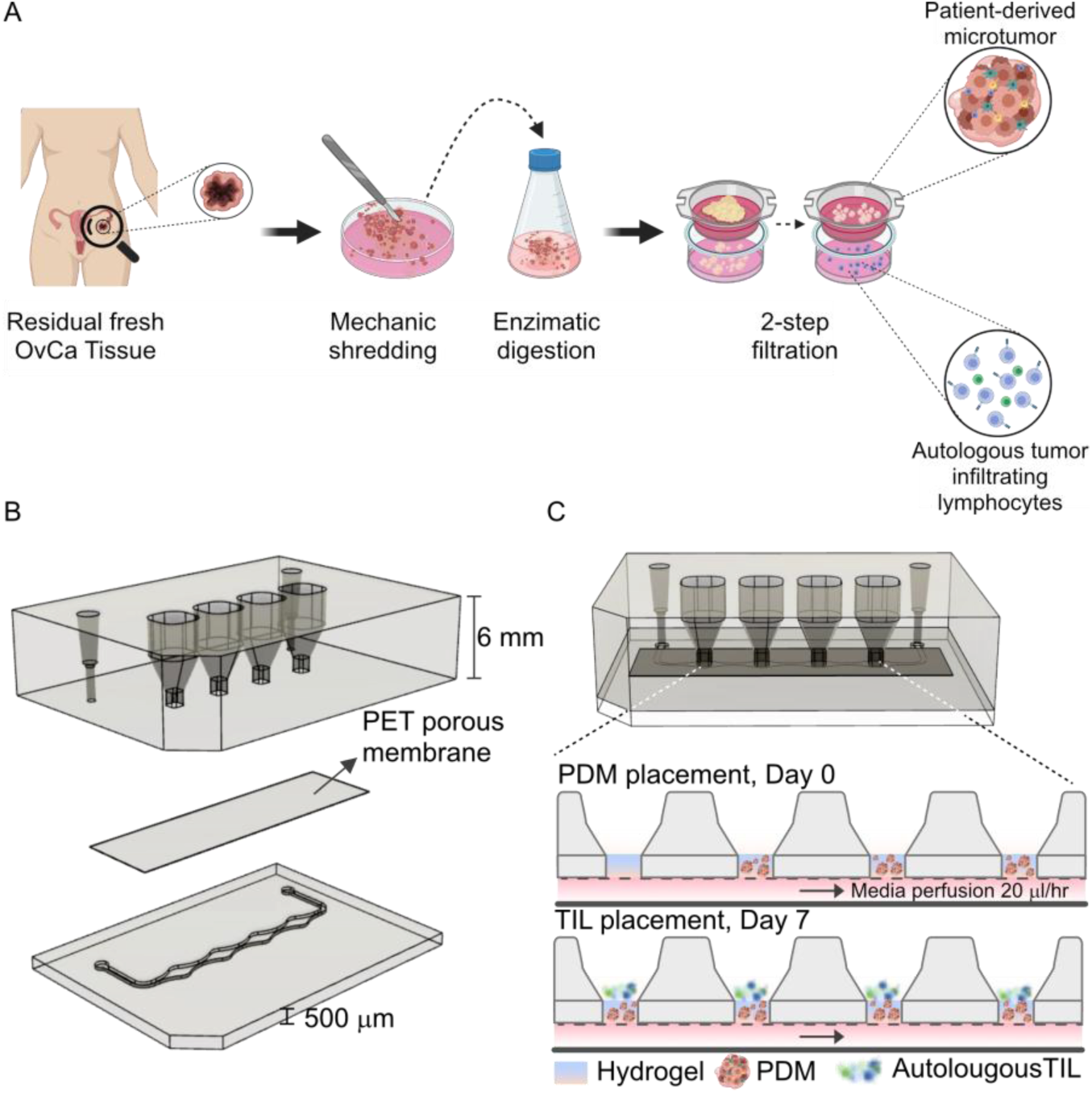
Schematic representation of the experimental workflow. (**A**) Isolation of patient-derived microtumors (PDM) and tumor-infiltrating lymphocytes (TILs) from fresh OvCa tissue specimens (created in Biorender). (**B**) Setup of the three-layer PDMS tumor-on-chip consisting of a 6 mm tissue layer with four serial tissue chambers, a 500 µm bottom layer containing the media channel and a porous PET membrane separating both layers. Layers were plasma bonded. (**C**) Depiction of sequential loading of OvCa PDM and TILs. On day 0, tissue chambers were loaded with OvCa PDM embedded in a hydrogel. Cell culture medium with or without paclitaxel was perfused at a constant flow rate of 20 µl/hr. On day 7, autologous TILs were added on top of the hydrogel, and the chips were perfused with media with or without ICI at the same flow rate until day 14.

### Viability measurement of OvCa PDM

To assess the viability of OvCa PDM in static and in microfluidic culture, live/dead-cell staining was performed using Calcein-AM™ (Invitrogen) live cell stain and SYTOX™ Orange nucleic acid dead stain (Invitrogen). Hoechst 33258 (Invitrogen) was added to visualize nuclei as described before (9, 10). Confocal z-stack images of stained PDM were taken by using a Zeiss CellObserver Z1 (Carl Zeiss). For perfused on-chip cultured OvCa PDM, chips were collected after 14 days.

OvCa PDM were stained on-chip using 100 µl of staining solution in a pipette tip placed in the inlet and an empty pipette tip placed in the outlet of the chips’ media channel. This was followed by an incubation period of 1 h at 37 °C. The staining solution is transported through the chip and into the empty pipette tip via hydostatic-driven flow (HDF). Z-stack images were taken using the Zeiss CellObserver Z1 (Carl Zeiss). Maximum intensity projection of the 3D z-stacks and image analysis was done using the ZEN software (Version 2.6). 3D surface masks for viable and dead stained cells were generated in the GFP and Cy3 channels, and fluorescence intensity sums of each channel were calculated and normalized to the total OvCa PDM volume.

### Co-culture of OvCa PDM and autologous TILs on-chip

The chips were flushed initially two times with 80 % ethanol (EtOH) and three times with DPBS. OvCa PDM were harvested by centrifugation (200x g, 5 min, RT), counted and resuspended in dextran hydrogel (Cellendes) supplemented with 0.5 mmol/L RGD peptide (Cellendes) (15). 50 PDM in 10 µl were seeded per chip, with 2 µl volumes per chamber. Prior to introducing the media into media channel, chambers were covered with PCR foil (Thermofisher Scientific), then flushed three times with PDM medium. This procedure prevents channel clogging by removing hydrogel deposits. After the hydrogel starts to crosslink, chips were incubated over night at 37 °C, 5 % CO_2_ in the incubator. The next day, chips were connected to an external syringe pumping system (LA-190, Landgraf Laborsysteme HLL GmbH). Perfusion was started with or without 25 µg/ml paclitaxel for six days with a flow-rate of 20 µl/h. On day 7, autologous TILs were administered for an additional seven days in the presence of the anti-PD-L1 antibody durvalumab (MedChemExpress) at different concentrations. Starting on day 4, flow-through was collected every 24 hours and stored at -80 °C / -20 °C until day 14.

To analyze the cytotoxic effect of the anti-PD-L1 antibody Durvalumab (MedChemExpress) with or without prior paclitaxel (Selleckchem) treatment and in combination with autologous TILs a caspase-cleaved cytokeratin 18 (ccCK18) ELISA (TecoMedical) was performed (16-18). Flow-through media from microfluidic chips was collected every 24 h starting on day 4 and stored at - 20 °C. After performing the assay, according to the manufacturer’s instructions, the assay was read using a Spark Multimode Plate Reader (Tecan).

### Bead-based cytokine and chemokine immunoassay analysis

For cytokine and chemokine analysis, flow-through was collected every 24 h starting on day 4 and stored at -80 °C. The LEGENDplex Human Proinflammatory Chemokine Panel 1 (13-plex) w/VbP (BioLegend) and LEGENDplex Human CD8/NK Cell Panel V02 (13-plex) w/VbP (BioLegend) assays were employed to examine the release of thirteen distinct chemokines, respectively. After sample processing according to manufacturer’s instructions, samples were measured using a BD FACS LSRFortessa (BD Biosciences) and analyzed using Qognit software (BioLegend).

### FACS analysis

For characterization of autologous TILs, up to 1x10^6 cells were harvested per sample, washed twice in PBS with centrifugation at 400x g for 5 min at 4 °C and resuspended in FACS staining buffer (0.2 mM EDTA, 10 FCS in PBS). To determine cell number and viability, cells were counted using a Nucleocounter (Chemotec). An unstained control was prepared for each TIL population. Prior to staining, Fc receptors were blocked with Human TruStain FcX™ (Biolegend) according to the manufacturer’s protocol. Cells were washed with FACS staining buffer (400x g, 5 min, 4 °C) and incubated with Zombie NIR™ Fixable Viability Kit (Biolegend), according to the manufacturer’s instructions to identify dead cells. Cells were washed with FACS staining buffer (400x g, 5 min, 4 °C) and antibodies for extracellular targets (Table S3) were applied and incubated for 30 min, at 4 °C in the dark. The cells were washed with FACS staining buffer (400x g, 5 min, 4 °C). Before antibodies for intracellular antigen staining were applied, cells were fixed and permeabilized with BD Pharmingen™ Transcription Factor Buffer Set (BD Bioscience) according to the manufacturer’s protocol. After two additional washing steps the cells were incubated with antibodies for 45 - 50 min at 4 °C in the dark. Subsequently, the cells were washed 2 - 3 times and then resuspended in 100-150 µl FACS staining buffer and analyzed using a BD FACS LSRFortessa (BD Biosciences). FlowJo™ software (BD Biosciences) was used to analyze the different cell populations. comprehensive overview of the applied antibodies can be found in Table S2.

### Statistical analysis

Statistical analysis was conducted using GraphPad Prism. Data are shown as mean ±SEM unless stated otherwise. Data sets were analyzed using a two-way ANOVA with multiple comparison test unless stated otherwise. Outliers were identified with the Iglewicz and Hoaglin’s robust test for multiple outliers applying a recommended Z-score of ≥3.5 (19). Spearman rank correlation coefficients were calculated for correlation analyses between TIL frequencies and functional tumor cell killing. For all analyses, p values < 0.05 were considered statistically significant. Semi-quantitative secretome rankings were assigned relative to the mediator-specific maximum TotalAUC across all OvCa samples and treatment conditions and were used for descriptive interpretation only. Clinical patient data and comparisons with ex vivo response profiles were analyzed descriptively.

## Results

### OvCa-on-Chip enables the creation of a spatially defined patient-derived TME using PDM for long-term ex vivo culture

To establish a customized microfluidic OvCa-on-chip that features tumor heterogeneity and central components of the TME, PDM and autologous TILs were isolated from residual OvCa primary tumor tissue within 24 hours after surgery (Fig. 1A), as previously described (9, 10, 13). The histopathological classification and corresponding FIGO stage of each tumor are listed in Table 1.

**Table 1:**
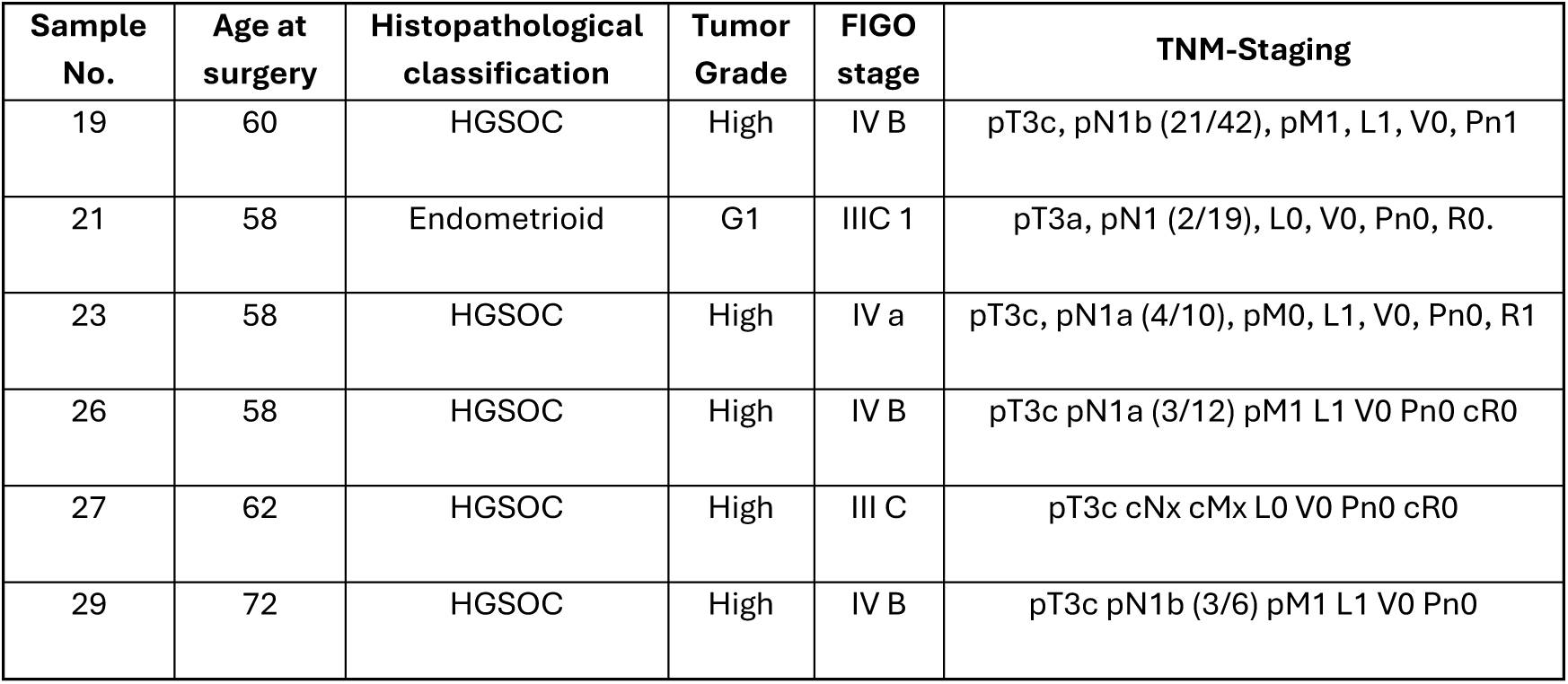
Clinical patient data from residual fresh OvCa tumor specimens included in this study for ex vivo on-chip cultures.

| Sample No. | Age at surgery | Histopathological classification | Tumor Grade | FIGO stage | TNM-Staging |
| --- | --- | --- | --- | --- | --- |
| 19 | 60 | HGSOC | High | IV B | pT3c, pN1b (21/42), pM1, L1, V0, Pn1 |
| 21 | 58 | Endometrioid | G1 | IIIC 1 | pT3a, pN1 (2/19), L0, V0, Pn0, R0. |
| 23 | 58 | HGSOC | High | IV a | pT3c, pN1a (4/10), pM0, L1, V0, Pn0, R1 |
| 26 | 58 | HGSOC | High | IV B | pT3c pN1a (3/12) pM1 L1 V0 Pn0 cR0 |
| 27 | 62 | HGSOC | High | III C | pT3c cNx cMx L0 V0 Pn0 cR0 |
| 29 | 72 | HGSOC | High | IV B | pT3c pN1b (3/6) pM1 L1 V0 Pn0 |

The microfluidic platform is composed of a three-layer polydimethylsiloxane (PDMS) chip comprising an upper tissue layer, a media channel layer at the bottom, and an intermediate porous PET membrane with a pore size of 3 µm (Fig. 1B). The tissue layer features four series-connected conical-shaped chambers, each designed to incorporate hydrogel-embedded OvCa PDM as spatially defined patient-derived TME units. The porous membrane physically separated the tissue chambers from the perfused media channel allowing diffusion of nutrients and treatment compounds, and the removal of waste products. Expanded autologous TILs were subsequently introduced into these PDM-containing compartments on day 7 to complement the native microenvironment with a patient-matched immune effector compartment and enable functional analysis of tumor-immune interactions under controlled flow conditions (Fig. 1C).

Histopathological features as well as the expression of tumor-specific markers and TME components of *n* = 6 patient-derived OvCa PDM were confirmed by HCE and immunohistochemical staining of FFPE (9, 10). OvCa PDM exhibited intertumoral heterogeneity of central TME components including collagen I, cancer-associated fibroblasts (FAPα), TILs (CD3^+^), and myeloid cells (CD68^+^). While OvCa21 expressed the full panel of markers shown in Fig. 2A, inter-donor variability was observed in tissue composition, with OvCa26 and OvCa27 lacking expression of some of these components. Importantly, we hereby confirmed the preservation of the native composition of the TME of all samples, underlining the value of these samples as physiologically relevant (9). Furthermore, the OvCa PDM isolated in this study exhibited robust viability across various tissue samples from different patients after isolation, with over 80% of live cells per PDM per model, as by Calcein-AM/Sytox orange live-dead cell staining (Fig. 2B-C, Fig. S2).

**Fig. 2:**
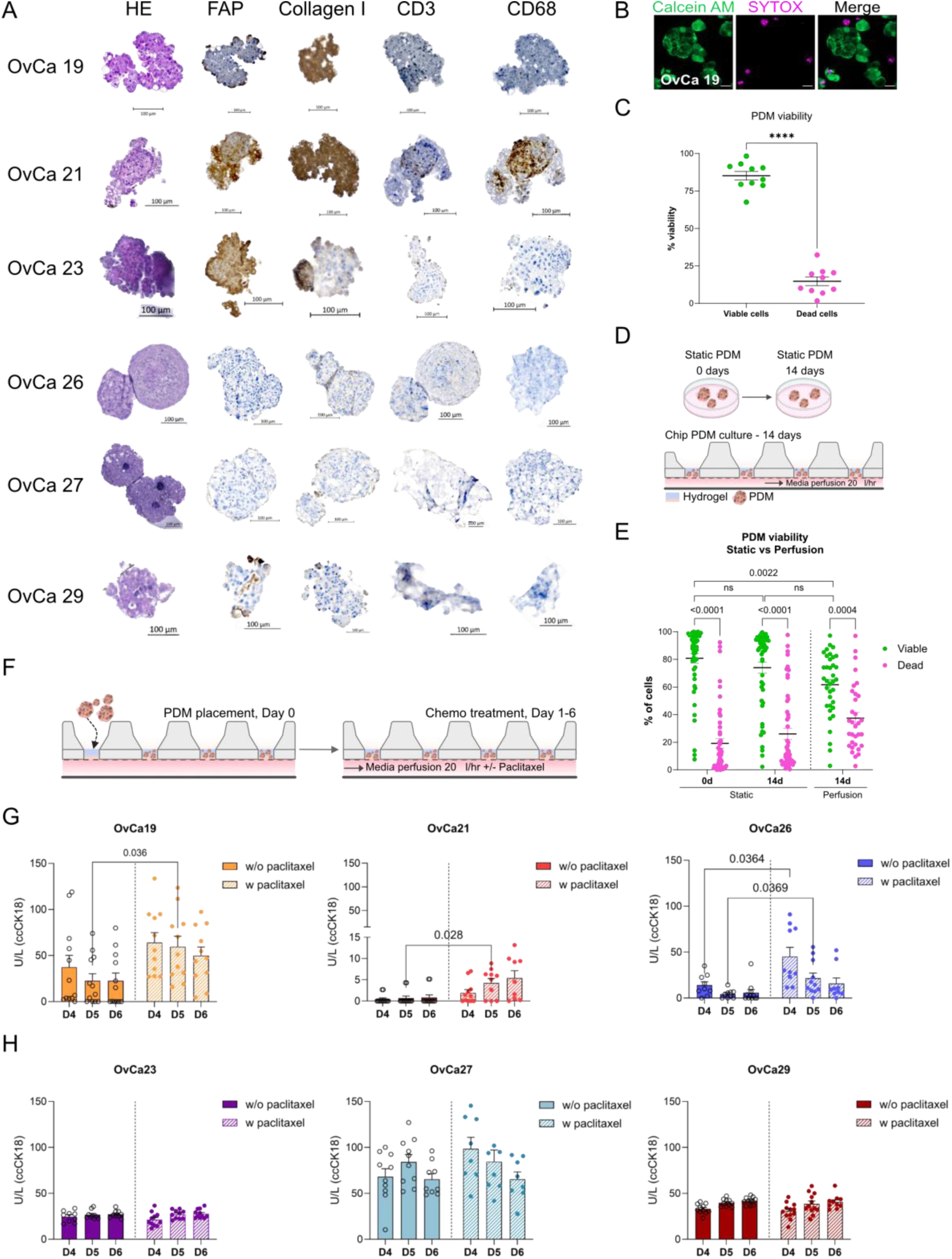
PDMs preserve features of native TME, maintain viability on-chip in perfused culture conditions and reflect heterogeneous, patient-specific responses to chemotherapy. (**A**) Representative immunohistochemical stainings for HCE, FAPα, collagen I, CD3 and CD68 on FFPE sections of OvCa PDM (shown for *n* = 6 different donors). (**B-C**) Viability assessment of OvCa PDM after isolation using Calcein-AM (live cells) and SytoxOrange (dead cells) (**B**) Exemplary 2D images from 3D projections of live/dead staining (shown OvCa19 PDM). (**C**) OvCa PDM show high viability after isolation from primary tumors. Viability was assessed by quantifying the percentage of live and dead cells from at least *n* = 6 single PDM per donor using image-based analysis (see Methods); each dot represents one donor (total *n* = 10 donors). Data are presented as mean ±SEM (Two-way ANOVA with Šídák’s multiple comparisons test). (**D-E**) OvCa PDM remain viable for up to 14 days in on-chip perfusion culture. (**D**) Experimental setup for viability comparison of PDM cultured for 14 days in static condition and perfused on-chip condition with *n* = 50 PDM per chip/static culture. (**E**) Live/dead cell staining was performed for n = 6 PDM donors. Percentage of live and dead cells from at least *n* = 6 single PDM per donor (static culture) and *n* = 3 chips per donor (on-chip perfusion culture) was quantified by image-based analysis (see Methods). Data are shown as mean ±SEM (Two-way ANOVA with Šídák’s multiple comparisons test). (**F**) Experimental workflow for the assessment of patient-specific chemotherapy response in perfused OvCa-on-chip cultures. Per donor *n* = 12 chips were seeded with 50 OvCa PDM each and perfused for 6 days with or without 25 µg/ml paclitaxel with a flow rate of 20 µl/hr. (**G-H**) OvCa-on-chip cultures reveal patient-specific responses to chemotherapy. Tumor cell death is monitored using caspase-cleaved cytokeratin 18 (ccCK18) ELISA from perfused media. Data are presented as mean ±SEM (Two-way ANOVA with Šídák’s multiple comparisons test). (**G**) Paclitaxel-sensitive donors (OvCa19, OvCa21 and OvCa26) show significant response towards chemotherapy. (**H**) Paclitaxel-resistant donors (OvCa23, OvCa27 and OvCa29) showed no response towards paclitaxel treatment. \**p* < 0.05; \*\**p* <0.01; \*\*\**p* <0.001; \*\*\*\**p* <0.0001; Scale bars (A) 100 µm, (B) 50 µm.

Long term culture of isolated PDM was achieved by seeding an average of 50 PDM per donor per microfluidic chip embedded in SG dextran-based hydrogel (Cellendes) supporting a 3D structure that mimics the functions of an extra-cellular matrix (ECM) (Fig.2D) (15). Loaded chips were connected to a syringe pump 24 hours after seeding to ensure continuous perfusion (flow rate: 20 µl/h) with media from the lower media channel. Medium replacement was performed every three days. PDM viability on the OvCa chip was assessed following 14 days of perfusion using live-dead cell staining with no significant reduction in cell viability compared to 14 days in conventional static culture (Figure 2E). These findings confirm the suitability of the microfluidic platform for long-term maintenance of viable patient-derived OvCa microtumors under dynamic culture conditions.

### OvCa PDM-on-chip enables dynamic monitoring of chemotherapy responses

To evaluate the platform for functional chemotherapy response profiling, OvCa PDM from *n* = 6 patients were treated on-chip with paclitaxel (PTX) under dynamic culture conditions (Fig. 2F). Tumor specific cell death was monitored daily by quantifying caspase-cleaved cytokeratin 18 (ccCK18) release in chip effluents as a proxy for epithelial tumor cell apoptosis. Treatment responses to PTX (25 µM for 6 days) showed patient heterogeneity in OvCa on-chip cultures. OvCa19, 21 and 26 displayed significant sensitivity to chemotherapy with increased ccCK18 release upon PTX exposure (Fig. 2G). In OvCa21, ccCK18 levels remained below the assay detection limit of 20 U/l in both untreated and PTX-treated cultures. Although this limits the interpretability of results from this model, a treatment-associated trend was nevertheless observed. In contrast, no significant paclitaxel-induced tumor cell death was detected in OvCa23, OvCa27, or OvCa29, indicating limited treatment sensitivity in these models during the analyzed culture period (Fig. 2H). These findings demonstrate that the OvCa-on-Chip system captures heterogeneous patient-specific chemotherapy responses under controlled perfusion conditions.

### On-chip profiling of sequential chemo-immunotherapy using longitudinal ccCK18 monitoring

As the established platform resolves donor-specific PTX responses and considering the general low response rates to ICIs in OvCa (20), we leveraged the same longitudinal readout to investigate patient-specific responses to sequential chemo-immunotherapy. OvCa PDM were cultured under continuous perfusion for 14 days and, where indicated, pretreated with PTX during the initial culture phase prior the addition of expanded autologous TILs on day 7. TILs were added either alone or in combination with PD-L1 blockade using DU at different concentrations (Fig. 3A). Tumor-specific cell death was monitored longitudinally by measuring ccCK18 release in chip effluents from day 4 onwards (Fig. 3B), as exemplified for OvCa19. Here, TIL addition induced detectable tumor cell death during the TIL-containing treatment phase from day 8 to day 14, as reflected by increased ccCK18 levels in the TIL treatment condition (d8-d14; -DU-PTX). Cumulative area under the curve (AUC) analysis of baseline-corrected ccCK18 values further confirmed TIL-mediated tumor cell death in this model (Fig. 3C). DU treatment alone did not enhance TIL-mediated killing, whereas PTX pretreatment enhanced tumor cell death following TIL addition, with only limited additional effects of subsequent DU treatment (combined therapy).

**Fig. 3:**
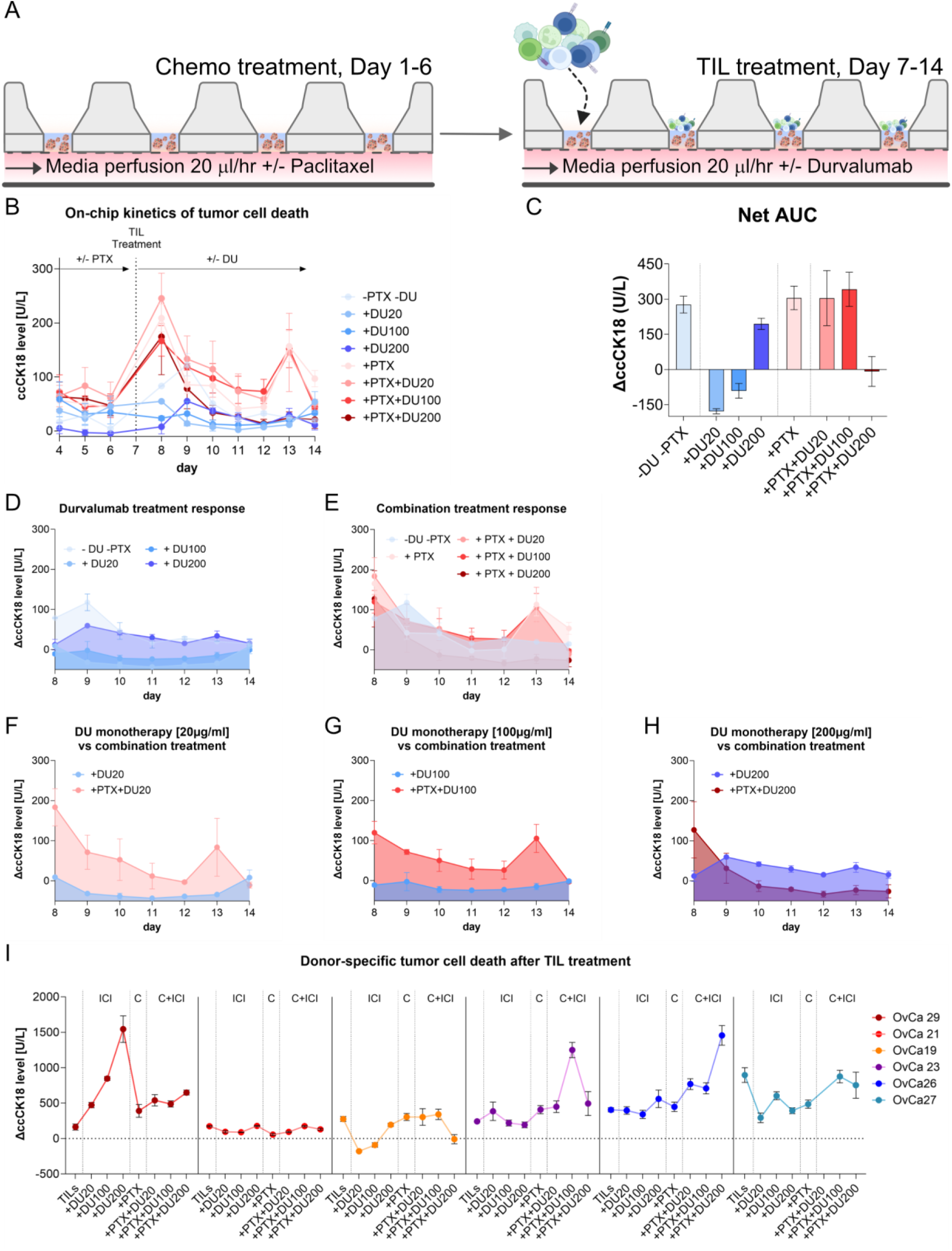
Functional response profiling of sequential chemo-immunotherapy in patient-derived OvCa-on-chip cultures. (**A**) Schematic overview of the experimental procedure. OvCa PDM (*n* = 50 per chip) were seeded in a dextran-based hydrogel and cultured under continuous perfusion from day 1 to 14. Where indicated, PDM were pretreated with 25 µg/ml paclitaxel (PTX) from day 0 to 6, prior to the addition of expanded autologous TILs (E:T 10:1) on day 7. Where indicated, PDM-TIL co-cultures were further treated with or without durvalumab (DU; 20, 100 or 200 µg/mL) until day 14. Chip effluents were collected every 24 h and analyzed for caspase-cleaved cytokeratin 18 (ccCK18) as a marker of epithelial tumor cell apoptosis. (**B-H**) Representative, functional response profiling in OvCa19 on-chip cultures. (**B**) Longitudinal ccCK18 kinetics of OvCa19 PDM-on-chip cultures from day 4 to day 14 across all treatment conditions. (**C**) Treatment-associated tumor cell death following TIL treatment. Values shown represent net area under curve (NetAUC) of ccCK18 values, corrected for the baseline (day 6), from day 8 to day 14. (**D–H**) Tumor cell death after TIL addition on day 7, as well as additional treatment effects of chemotherapy pretreatment with or without PD-L1 blockade demonstrated as kinetic data (day 8-14, values corrected for baseline day 6). (**D**) Treatment response to anti-PD-L1 therapy compared to TIL-treatment. (**E**) Treatment response to sequential chemo-immunotherapy (**F–H**) The impact of PTX pretreatment on the efficacy of PD-L1 blockade. Direct comparison of DU treatment with the corresponding combination PTX+DU treatment at DU20, DU100 and DU200, respectively. (**I**) Visualisation of patient-specific treatment responses using NetAUC (baseline-corrected ccCK18 values, day 8 to day 14). Data are shown as mean values ±SEM from *n* = 3 chips.

### Time-resolved ccCK18 kinetics distinguish TIL-, PTX- and DU-associated effects in OvCa13

To further dissect treatment-specific effects over time, we compared baseline-corrected ccCK18 kinetics between the respective treatment conditions (Fig. 3D-H). While TIL treatment alone induced a transient increase in ccCK18 release, DU-treated conditions largely remained within similar or lower range, indicating that PD-L1 blockade alone had no additional effect on TIL-mediated killing in OvCa19 (Fig. 3D). In contrast, PTX pretreatment altered TIL tumor cell death kinetics. Compared with TIL treatment alone, PTX-pretreated conditions showed increased ccCK18 release immediately after TIL addition (Fig. 3E). Similarly, this effect was also observed between days 12 and 14 compared to single TIL therapy (-PTX-DU). The combinations of DU and PTX did not uniformly increase ccCK18 release across the full observation period, indicating that the major increase in tumor cell death was mainly associated with PTX pretreatment. Direct comparison of DU monotherapy with the corresponding PTX-DU combination further supported this finding (Fig. 3F-H). Together, these time-resolved comparisons indicate that tumor cell death in OvCa19 was primarily driven by TIL addition after day 7 and enhanced by prior paclitaxel exposure. PD-L1 blockade with durvalumab induced only modest and non-uniform additional effects.

### Patient-derived OvCa-on-chip systems show distinct response patterns to TILs, PD-L1 blockade and sequential chemo-immunotherapy

Across additional OvCa models, NetAUC values (Fig. 3I, Supplementary Fig. S4A-B) revealed pronounced donor-specific treatment patterns, which were further summarized by ranking treatment responses assigned by calculating fold changes of matched treatments in Table S4. This integrated analysis distinguished models with high baseline TIL-mediated killing from those showing treatment-specific enhancement. OvCa23 and OvCa26 showed the clearest increase in tumor cell death following sequential PTX+DU treatment with OvCa23 displaying the strongest relative combination effect and OvCa26 showing a more sustained PTX+DU-associated response (Fig. 3I, Supplementary Fig. S4A-B). In contrast, OvCa29 responded predominantly to TIL+DU treatment without PTX pretreatment in a dose-dependent manner (Fig. 3I, Supplementary Fig. S4A-B). OvCa27 showed high TIL-only ccCK18 release but no consistent additional benefit from PTX or DU, whereas OvCa21 showed only limited treatment-induced ccCK18 responses (Fig. 3I). Time-resolved comparison of DU-treated conditions further demonstrated that PD-L1 blockade alone had donor-dependent effects. In OvCa29, TIL+DU treatment was associated with a sustained increase in ccCK18 release. This increase was dose-dependent. In contrast, DU effects were weaker or more transient in OvCa21, OvCa23, and OvCa27 (Supplementary Fig. S4C). When comparing the effects of PTX pretreatment, OvCa23 displayed a sharp transient peak, while OvCa26 showed a more sustained response pattern (Supplementary Fig. S4D). Direct comparison of DU monotherapy with its respective PTX+DU combinations revealed that PTX pretreatment improved DU-treatment notably in OvCa23 and OvCa26. However, chemotherapy pretreatment was not uniformly beneficial across patient models (Supplementary Fig. S4E–G). These data indicate that the system can capture the kinetics of patient-specific sensitivity to TIL-mediated killing, PD-L1 dose-dependent blockade, and sequential chemo-immunotherapy treatment.

### Longitudinal secretome profiling reveals treatment-dependent TIL effector activity

To complement the ccCK18-based tumor cell death readout, cytokines and chemokines release reflecting cytotoxic immune activity and inflammatory TME signaling were quantified. In the example OvCa19, TIL addition on day 7 induced a temporally structured effector response, with early IFNy secretion followed by delayed granzyme A release and subsequent granulysin and perforin secretion (Fig. 4A; additional data of all OvCa donors in Supplementary Fig. S5). Higher DU concentrations enhanced granzyme A, granulysin, and perforin levels compared with TIL treatment alone, whereas PTX pretreatment broadened and prolonged cytotoxic mediator secretion across several conditions. Overall, PTX-pretreated cultures tended to show a modest delay in peak secretion, indicating that chemotherapy pretreatment altered the kinetics of the subsequent TIL effector response. Chemokine release followed a distinct but related kinetic pattern. CCL5 secretion increased rapidly after TIL addition across several treatment conditions (Figure 4B) and remained detectable throughout the TIL-containing phase (day 8 to day 14), consistent with an activated lymphocyte-associated chemokine response.

**Fig. 4:**
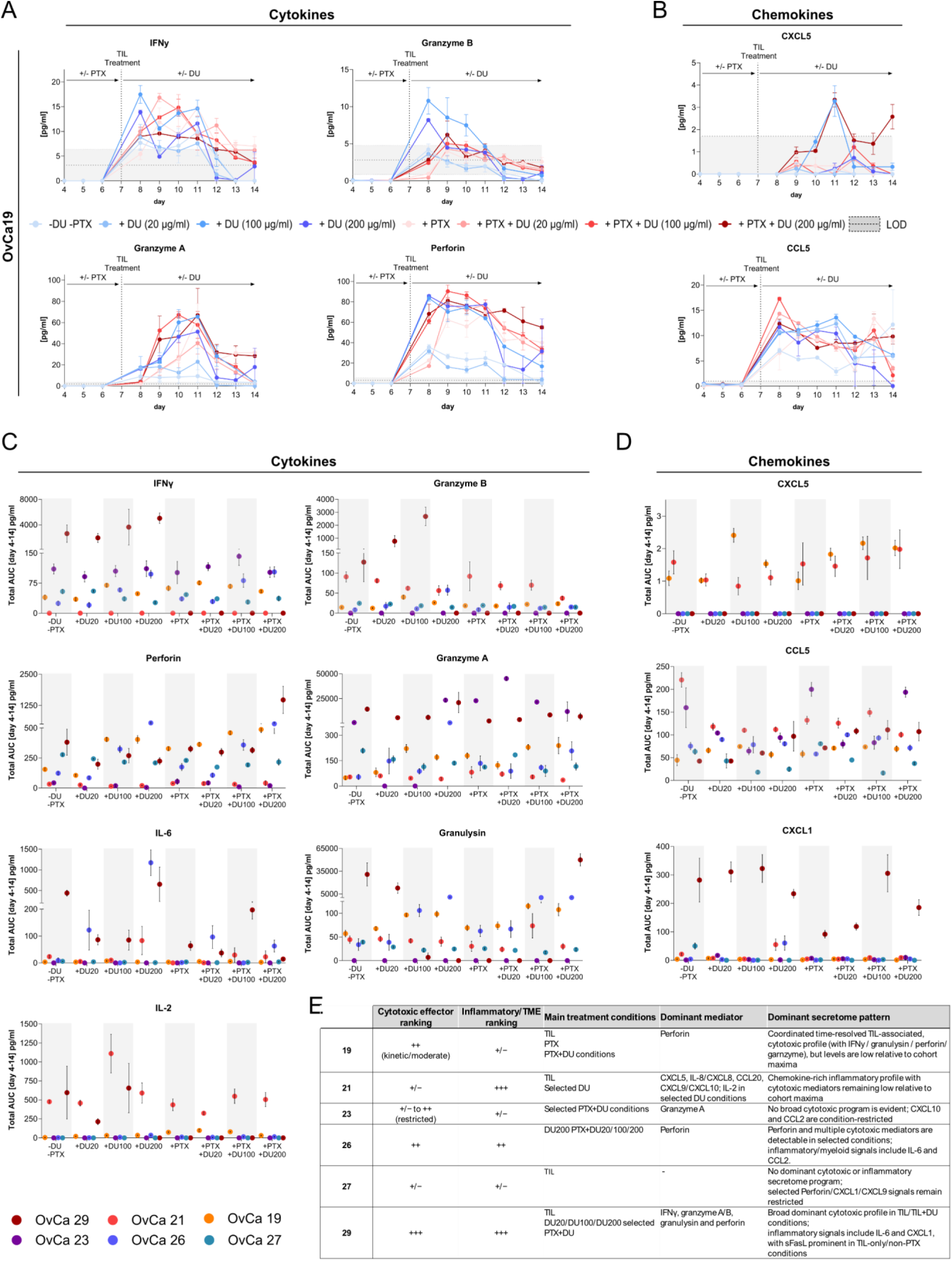
Longitudinal secretome profiling reveals donor- and treatment-dependent patterns. (**A**) Representative kinetics of cytotoxic TIL effector mediators in OvCa19 PDM-on-chip cultures. IFNγ, granzyme A, granzyme B, granulysin and perforin were quantified in chip effluents collected from day 4 to day 14. (**B**) Representative chemokine secretion kinetics of CXCL5 and CCL5 in OvCa19 PDM-on-chip cultures across the indicated treatment conditions in OvCa19 PDM-on-chip cultures. (**C-D**) Cumulative cytokine and chemokine profiles across patient-derived OvCa models and treatment conditions. Total AUC values were calculated from effluent cytokine/chemokine concentrations measured from day 4 to day 14. Each color represents one donor. (**E**) Semi-quantitative summary of dominant secretome patterns across OvCa-on-chip cultures. Cytokine and chemokine profiles were summarized based on TotalAUC values calculated from day 4 to day 14 and ranked in three categories: high/dominant (+++), moderate/detectable (++), and low/restricted (+/−). Rankings were assigned relative to the mediator-specific maximum across all models and treatment conditions and were used for descriptive interpretation only. The table summarizes cytotoxic effector activity, inflammatory/TME-associated signaling, dominant mediators, and treatment conditions associated with the most pronounced secretome responses.

### Multi-analyte secretome profiling captures donor-specific immune activation and inflammatory TME signaling

To compare the cumulative secretion of cytokines and chemokines across treatment conditions and patients, the TotalAUC were calculated from measured cytokine/chemokine values of day 4 to 14. Secretome profiles were highly treatment- and donor-dependent, indicating that treatment-induced tumor cell death was accompanied by heterogeneous immune and inflammatory response states rather than a uniform secretory program (Fig. 4C; Supplementary Fig. S6). To facilitate the interpretation of secretome profiles, dominant secretome patterns were summarized using a semi-quantitative ranking of cytotoxic and inflammatory mediators (Fig. 4E). In OvCa29, which significantly responded to DU-monotherapies (Fig. 3I, Supplementary Table S4), TIL treatment induced high levels of IFNγ, granzyme A, perforin and granulysin (Fig. 4C-E). The addition of DU further boosted selected cytotoxic mediators, strikingly IFNy and granzyme B. However, this DU-associated pattern was abolished in PTX-pretreated conditions. Granzyme A levels remained high across most of the treatments, suggesting a strong TIL-associated baseline effector state in OvCa29 rather than a treatment-specific response (Fig. 4C-E, Supplementary Fig. S5-6). By contrast, OvCa19 showed a more moderate, but superior retention cytokine secretion of cytotoxic mediators (Figure 4C-E, Supplementary Fig. S5-6), in line with the TIL-mediated tumor cell death observed in the kinetic ccCK18 analysis (Figure 3B, Supplementary Fig. S4). OvCa 26 displayed treatment-enhanced cytotoxic mediator release in selected DU- and PTX+DU combination treatments, indicative of treatment-dependent modulation of cytotoxic TIL activity (Fig. 3I, 4C-E, Supplementary Fig. S4). OvCa23, however, showed a more restricted and transient profile, with selected mediator signals but no broad cytotoxic effector pattern comparable to OvCa19, OvCa26 or OvCa29 (Fig. 4C-E, Supplementary Fig. S5-6). Thus, cytotoxic mediators complemented ccCK18-based tumor cell death quantification by providing information on the extent of TIL effector activation. However, these mediators did not uniformly mirror ccCK18 release across all donors, indicating that immune activation and measurable tumor cell death were only partially coupled.

Inflammatory and TME-associated mediators such as IL-6, CXCL1, CXCL5, displayed donor- and treatment-dependent patterns (Fig. 4C-D). CXCL1 displayed elevated levels mainly in OvCa29, particularly in TIL and DU-treated containing conditions (Fig. 4D). IL-6 secretion was increased in OvCa29 and OvCa26, particularly in high doses of DU treatment (Fig. 4C). High CXCL5 AUCs were mostly restricted to OvCa21 and OvCa19, suggesting a donor-specific inflammatory TME state. By contrast, CCL5 was among the most consistent chemokine readouts detected across models and treatment conditions, supporting its use as a marker of lymphocyte-associated chemokine activity (Fig. 4D).

Collectively, these results show that OvCa-on-chip system provide cytokine and chemokine profiling that allow to resolve treatment-specific immune activation and patient-dependent inflammatory TME responses beyond ccCK18-based tumor cell death quantification.

### Association of TIL subpopulations with TIL-mediated tumor cell killing

To relate functional treatment responses to the composition of the autologous immune compartment, expanded TILs products used for on-chip co-culture were characterized by multiplex flow cytometry. TIL composition varied highly across the different OvCa patients (Fig. S3B–E). While OvCa29, OvCa21 and OvCa19 were characterized by higher CD3^+^ T cell frequencies, OvCa23, OvCa26 and OvCa27 contained substantially lower proportions of CD3^+^ T cells (Fig. S3B). Within the CD3^+^ T cell fraction we found high proportion of cytotoxic CD8^+^ T cells in OvCa26 (68.3%), whereas modest levels were found in OvCa19 (33.1%) and OvCa29 (26.7%) (Fig. S3C). In contrast, OvCa21 and OvCa23 T cells were predominantly enriched for CD4⁺ T helper cells (Fig. S3D), indicating substantial interpatient heterogeneity in TIL product composition.

We next assessed whether distinct specific TIL phenotypes were associated with functional treatment responses by correlating TIL subset frequencies with the ccCK18 NetAUC across the respective treatment conditions. Several TIL subsets were identified that positively correlated with enhanced tumor cell killing under treatment naïve conditions (Fig. 5A). Specifically, a significant positive correlation was observed between TIL-mediated tumor cell killing and the frequency of naïve CD4^+^ TILs (Spearman’s *r* = 0.94, *p* = 0.017), as well as CD4^+^PD-1^+^ TILs (Spearman’s *r* = 0.94, *p* = 0.017). Checkpoint-defined CD4⁺ T cell subsets showed additional associations with TIL-mediated killing. Frequencies of CD4⁺PD-1⁺TIM-3⁺ and CD4⁺PD-1⁺TIM-3⁻ TILs positively correlated with TIL-mediated tumor cell killing (Spearman’s *r* = 0.89, *p* = 0.033 and *r* = 1.00, *p* = 0.003, respectively), whereas CD4⁺PD-1⁻TIM-3⁻ TILs showed a negative correlation (Spearman’s *r* = −0.94, *p* = 0.017). Although limited to a small number of patient samples these correlations suggest that PD-1-expressing CD4⁺ TIL subsets, rather than checkpoint-negative CD4⁺ populations, were associated with functional antitumor activity in this assay. Within CD8 TIL subsets, a positive association was observed for CD8⁺CD69⁻CD103⁻ TILs (Spearman’s *r* = 0.94, *p* = 0.033). In contrast, the frequency of CD8⁺CD69⁺CD103⁻ TILs showed a negative correlation (Spearman’s *r* = −0.94, *p* = 0.017).

**Fig. 5:**
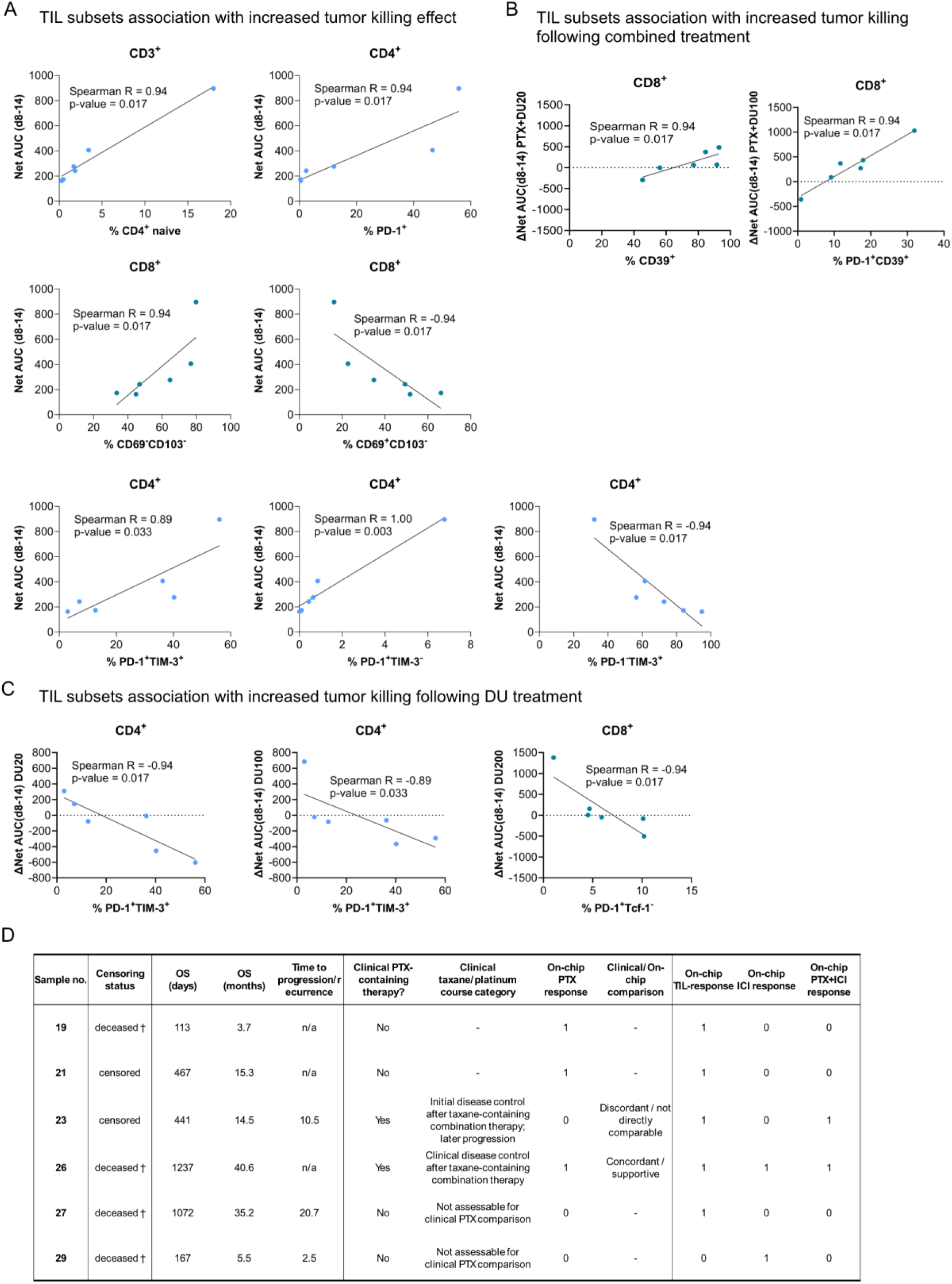
Exploratory correlation analysis of TIL composition and treatment-specific on-chip responses. Frequencies of selected CD4⁺ and CD8⁺ TIL subsets were correlated with functional tumor cell killing across patient-derived OvCa models. (**A**) Spearman correlation of expanded autologous TIL subsets and ccCK18 NetAUC values (day 8 – 14) during TIL-mediated tumor cell killing. (**B-C**) DU- or combination-associated effects. ΔNetAUC values were calculated by subtracting the corresponding matched reference condition, thereby capturing the additional treatment-induced tumor cell death beyond baseline TIL-mediated killing. (**B**) Spearman correlation of TIL subsets and anti-PD-L1 response. (**C**) Spearman correlation of TIL subsets and sequential combination treatment of PTX and DU. (**D**) Clinical patient data wiht overall survival (OS), time to recurrence and response of PDM per patient in relation to PTX, ICI and combined treatments on chip (“0” = no treatment response; “1” = treatment response).

### TIL subpopulations associated with durvalumab and sequential chemo-immunotherapy-induced therapy response

Treatment-specific correlations further indicated that the relationship between TIL phenotype and tumor cell killing depended on the therapeutic context. Following DU treatment, CD4⁺PD-1⁺TIM-3⁻ TILs correlated negatively with tumor cell killing at DU20, while CD4⁺PD-1⁺TIM-3⁺ TILs showed a negative association at DU100. In addition, CD8⁺PD-1⁺Tcf1⁻ TILs negatively correlated with tumor cell killing under high-dose DU treatment, consistent with a terminally exhausted phenotype being less favorable for PD-L1 blockade-associated responses (Fig.5B).

Under sequential PTX+DU treatment, CD8⁺CD39⁺ TILs and CD8⁺PD-1⁺CD39⁺ TILs positively correlated with tumor cell killing at PTX+DU20 and PTX+DU100, respectively (Fig.5C). Given that CD39 expression has been linked to tumor antigen-experienced CD8⁺ TILs, these findings suggest that tumor-reactive CD8⁺ T-cell subsets may contribute to functional responses in selected chemo-immunotherapeutic settings. Notably, the abundance of CD8+PD-1+Tcf1− TILs showed a significant negative correlation with tumor cell killing under durvalumab treatment (200 µg/mL) (Spearman’s *r* = −0.94, *p* = 0.017).

Taken together, these exploratory correlations indicate that tumor cell killing in the OvCa-on-chip platform is associated with distinct TIL phenotypes in a treatment-dependent manner and supports the use of integrated phenotypic and functional profiling to identify immune features linked to patient-specific responses.

### Evaluation of available clinical patient data and OvCa on-chip response profiles

To contextualize the functional on-chip response patterns, available clinical follow-up data were summarized for the corresponding patient-derived OvCa models (Fig. 5D). The cohort included clinically heterogeneous cases with marked differences in overall survival/follow-up time and time to progression or recurrence. Clinical taxane/platinum-containing treatment information was available for OvCa23 and OvCa26. OvCa26 showed clinical disease control after taxane-containing combination therapy and a concordant on-chip PTX response, supporting the relevance of the *ex vivo* chemotherapy readout in this model. In contrast, OvCa23 showed initial clinical disease control followed by later progression, while the on-chip assay indicated limited PTX sensitivity. These observations suggest that this patient may benefit from a combined chemotherapy regimen targeting both cytoskeletal dynamics and DNA cross-linking. Although not addressed in the present study, this standard-of-care combination could be readily incorporated into the longitudinal experimental frameworks that the OvCa-on-chip platform enables. Nonetheless, we observed an enhanced response under sequential PTX+ICI treatment, highlighting a more complex interplay between chemotherapy and immune checkpoint inhibition. The remaining patients have not received PTX-based adjuvant therapy, limiting the number of possible comparisons.

## Discussion

Although microfluidic cancer models have advanced considerably across several tumor entities, OvCa-specific platforms for patient-specific therapy response profiling remain limited (21-24). Existing OvCa-on-chip approaches have largely focused on diagnostic applications (reviewed in: (25)) or metastatic dissemination, while treatment studies often rely on established cell lines or patient-derived xenograft (PDX), which either incompletely reflect HGSOC heterogeneity, lack an autologous immune component or require time-consuming *in vivo* expansion (26-32). In this study, we developed an advanced patient-derived OvCa-on-chip platform that integrates patient-derived OvCa microtumors with sequential integration of autologous TILs in a microfluidic chip. This approach enabled longitudinal functional profiling of different single and combination treatment regimens of chemotherapy and ICIs.

A central feature of this platform is the combination of PDM with a dynamic perfusion. OvCa PDM have been extensively characterized previously and were shown to recapitulate subtype-specific characteristics and molecular properties of the original primary tumors (9). Together with continuous perfusion, this enabled sustained culture and controlled exposure to TIL-treatment, chemotherapy, single ICI or sequential chemo-immunotherapy within 14-day culture period. By this approach, key limitations of current preclinical models, which often fail to replicate the patient-specific tumor heterogeneity, therapy resistance, the immune contexture and the dynamic interplay between tumor and immune compartment (7, 8), especially when influenced by treatment, are directly addressed. This longitudinal design was particularly important for dissecting the treatment-induced tumor cell death and dynamic cytokine secretion patterns, thereby allowing simultaneous evaluation of functional tumor and immune responses.

Functionally, the platform captured interpatient variability in PTX sensitivity, including both sensitive and poorly responsive models and further distinguished early chemotherapy-associated effects from subsequent TIL- and PD-L1 blockade-associated responses. In the sequential chemo-immunotherapy setting, responses to TIL addition, PD-L1 blockade, and PTX pretreatment were also highly patient-specific. Notably, PTX pretreatment enhanced subsequent TIL- or DU-associated tumor cell killing in selected models, whereas in others PD-L1 blockade was more effective without prior chemotherapy exposure (Supplementary Table S4). Across the cohort, we observed treatment and donor-specific responses to treatments, underscoring the necessity of personalized functional approaches. Combining tumor cell death analyses with secretome profiling provided a more comprehensive view of tumor-immune responses. While IFNγ, granzyme A/B, granulysin and perforin reflected cytotoxic effector activity, additional mediators such as IL-2, IL-6, IL-8, IL10, CXCL1, CXCL5 and CCL20 showed more restricted donor- and treatment-dependent patterns. Proliferative signaling as reflected by IL-2 release was restricted to selected OvCa21 and OvCa29 conditions, suggesting model-specific T-cell proliferation, whereas IL-8/CXCL8, CXCL5, IL-6 and CXCL1 reflected broader inflammatory TME signaling. In OvCa21, high IL-8/CXCL8, CCL20 and CXCL5 levels pointed to a pronounced inflammatory chemokine signaling that is further enhanced after TIL addition. The secretion might be linked to high abundance of macrophages in the PDM (CD68^+^ IHC staining; Figure 2A), suggesting a myeloid-associated inflammatory TME state. In contrast, OvCa29 showed a broader inflammatory and cytotoxic profile, combining DU-associated tumor cell death with increased cytotoxic mediators and elevated IL-6 and CXCL1 in selected conditions. These mediator profiles were strongly donor- and treatment-dependent and did not consistently mirror ccCK18 release, which only reflects caspase-dependent apoptotic cell death. However, the combined analysis of apoptotic tumor cell death and multi-analyte secretome profiles provided a more comprehensive view of the occuring tumor–immune responses than either readout alone.

Importantly, the system enabled direct linking of treatment outcomes to the phenotypic composition of autologous immune cells. Distinct TIL subpopulations were associated with differential tumor cell killing depending on the treatment context, suggesting that on-chip functional responses were influenced by the specific composition of the TILs. In particular, PD-1 expressing CD4⁺ TILs were associated with baseline cytotoxic activity (33), suggesting that activated helper T-cell compartments can contribute to antitumor responses. However, T helper cells were not fully dissected to attribute this finding to a particular T helper cell subset. In contrast, terminally exhausted CD8⁺PD-1⁺Tcf1⁻ T cells correlated negatively with DU responses, consistent with their limited proliferative capacity and reduced responsiveness to checkpoint blockade (34-36). Conversely, CD8⁺CD39⁺ TILs were associated with enhanced tumor cell killing under sequential PTX and DU treatment, consistent with the reported link between CD39 expression and tumor-antigen-experienced CD8⁺ TILs (37). Positive association of CD8⁺CD69⁻CD103⁻ TILs hints towards non-resident or less tissue-adapted CD8⁺ T-cell populations (38) may retain relevant effector capacity in this *ex vivo* setting. This suggests that phenotypic markers of tissue adaptation alone are insufficient to predict functional efficacy in expanded autologous TIL products, particularly under *ex vivo* culture conditions. Taken together, we demonstrate a comprehensive characterization of TIL subpopulations applied within the OvCa-on-chip platform and establish a clear association between distinct TIL phenotypic states and corresponding treatment responses.

Several limitations should be acknowledged. This study is reflected with the availability of primary patient material, which constrained the scale of downstream analyses. Additionally, limited PDM yield from some of the primary tissue specimen, the tumor geography and quality of the sample obtained, restricts a full description and analyses. With the small patient cohort of this study, correlation analyses should therefore be considered exploratory. A further limitation is that the platform does not model de novo recruitment of immune cells from the circulation. Instead, it focuses on preserving key local TME features of patient-derived OvCa microtumors and functionally integrating patient-matched autologous TILs to study tumor-immune interactions within a controlled *ex vivo* setting. Expanded TILs also do not fully reproduce the native immune landscape of the original tumor, as *ex vivo* expansion may alter cell frequencies and clonal composition. Finally, ccCK18 specifically captures apoptotic epithelial tumor cell death, and additional readouts such as total CK18/M65, LDH release, live imaging, or spatial immune-cell tracking may help assess non-apoptotic tumor cell death in future studies. Future work should also extend the platform to additional clinically relevant treatment combinations, including emerging chemo-immunotherapy regimens and alternative immune checkpoint targets. Nevertheless, the workflow established here has clear translational potential as a framework for functional therapy response profiling. Rapid isolation of OvCa PDM and autologous TILs from patient tumor tissue, followed by controlled perfusion culture and sequential on-chip treatment within 14 days, enables patient-specific response data to be generated within clinically relevant timeframes. By combining longitudinal tumor cell death monitoring, secretome profiling and correlative TIL phenotype analysis, the platform provides complementary functional readouts. Future studies in larger prospective cohorts will be required to determine whether these response profiles correlate with clinical outcomes and can support patient stratification.

## Conclusion

The results of this study establish a novel, versatile, patient-derived OvCa-on-chip platform for the longitudinal functional profiling of chemotherapy, TIL-mediated tumor cell killing, PD-L1 blockade, and sequential chemo-immunotherapy. The system captured pronounced interpatient heterogeneity in therapeutic sensitivity and tumor-immune response patterns by integrating dynamic tumor cell death monitoring, multi-analyte secretome profiling, and correlative phenotyping of autologous TIL subsets. Beyond assessing treatment responses, the platform provides a controlled *ex vivo* framework to investigate immune functional states, treatment-associated inflammatory remodeling, and potential markers of response or resistance. Future studies in larger, prospectively collected patient cohorts, as well as correlation with clinical outcomes, will be essential to define the platform’s predictive value and support its further development toward functional precision oncology in OvCa.

## List of Abbreviation

2D: two-dimensional
3D: three-dimensional
AUC: area under the curve
CCK18: caspase-cleaved cytokeratin 18
CCL: C-C motif chemokine ligand
CIVM: complex in vitro model
CO2: carbon dioxide
CTCs: circulating tumor cells
CXCL: C-X-C motif chemokine ligand
DAB: 3,3′-diaminobenzidine
DMSO: dimethyl sulfoxide
DPBS: Dulbecco’s Phosphate-Buffered Saline
DU: durvalumab
DU20: durvalumab 20 µg/ml
DU100: durvalumab 100 µg/ml
DU200: durvalumab 200 µg/ml
ECM: extra-cellular matrix
EDTA: ethylenediaminetetraacetic acid
ELISA: enzyme-linked immunosorbent assay
EpCam: epithelial cell adhesion molecule
E:T: effector:target
EtOH: ethanol
FACS: fluorescence-activated cell sorting
FAPα: fibroblast activation protein alpha
FCS: fetal calf serum
FFPE: formalin-fixed paraffin-embedded
FIGO: International Federation of Gynecology and Obstetrics
HCE: hematoxylin and eosin
HDF: hydrostatic-driven flow
HGSOC: high-grade serous ovarian carcinoma
hr: hour
ICB: immune checkpoint blockade
ICI: immune checkpoint inhibition
IFNγ: interferon gamma
IL: interleukin
L: liter
min: minute
ml: milliliter
mm: millimeter
mmol: millimole
NK: cells natural killer cells
OvCa: ovarian cancer
PBS: phosphate-buffered saline
PCR: polymerase chain reaction
PD-1: programmed cell death protein 1
PD-L1: programmed death-ligand 1
PDM: patient-derived microtumors
PDMS: polydimethylsiloxane
PECVD: plasma-enhanced chemical vapor deposition
PET: polyethylene terephthalate
RTU: ready to use
PDM: patient-derived microtumors
PDX: patient-derived xenograf
PTX: paclitaxel
RT: room temperature
SG: slow gelling
Tcf1: T cell factor 1
TILs: tumor-infiltrating lymphocytes
TIM-3: T-cell immunoglobulin and mucin-domain containing protein
TME: tumor microenvironment
ToC: tumor-on-chip
Trm: tissue-resident memory cells
U: units
µg: microgram
µl: microliter
µM: micromolar
VbP: V-bottom plate
°C: degrees Celsius

## Declarations

### Ethics approval and consent to participate

Ovarian tumor tissue was collected from six OvCa patients undergoing surgery at the Center for Women’s Health, University Hospital Tübingen. Tumors were categorized according to International Federation of Gynecology and Obstetrics (FIGO) grading system. Written informed consent was obtained and the project was approved by the ethics committee at the Medical Faculty Tuebingen (703/2019BO2).

### Consent for publication

Not applicable.

### Availability of data and materials

All data generated or analyzed during this study are included in this published article [and its supplementary information files].

### Competing interests

TE: Abbvie, Astra Zeneca, Daiichi Sankyo, GSK, Gilead, Novartis, Eli Lilly, Pfizer, Pierre Fabre, MSD, Stemline AH: Speakers Bureau: Roche, Novartis, Lilly, MSD, AstraZeneca, DaichiiSankyo, Seagen, GSK, ExactScience, Gilead, Menarini Stemline, Pfizer, Eisai, Veracyte; Honoraria: Roche, Novartis, Lilly, MSD, AstraZeneca, Agendia, Seagen, GSK, ExactScience, Riemser, Teva, Onkowissen, Gilead, Menarini Stemline, Pfizer, Amgen, Pierre Fabre, Eisai, DaichiiSankyo, Thieme, Veracyte, Springer; Consulting or advisory role: Roche, Novartis, MSD, Agendia, AstraZeneca, GSK, ExactScience, Riemser, Teva, Onkowissen, Lilly, Gilead, Menarini Stemline, Pfizer, Amgen, Pierre Fabre, DaichiiSankyo; Travel support: Roche, Novartis, Lilly, AstraZeneca, GSK, ExactScience, Gilead, Menarini Stemline, Pfizer, DaichiiSankyo, Receipt of grants/research supports: ExactScience, Veracyte SP, NA, AR, EW, TS, JR, TIM, LC, AK, KSL, CS, PL: The authors declare that they have no competing interests.

### Funding

This work was supported by funding from the Wellcome Leap HOPE Program.

### Authors’ contributions

**SP:** Conceptualization; data curation; formal analysis; investigation; methodology; writing – original draft; writing – review and editing. **NA:** Conceptualization; data curation; formal analysis; investigation; writing – original draft; writing – review and editing. **EW:** investigation; writing – review and editing. **TS:** investigation; writing – review and editing. **JR:** investigation; writing – review and editing. **TIM:** methodology, writing – review and editing. **LC:** methodology, writing – review and editing. **TE:** data curation; writing – review and editing. **AH:** data curation; writing – review and editing. **AK:** data curation; writing – review and editing. **SYB:** data curation; writing – review and editing. **KSL:** writing – review and editing. **AR:** writing – original draft; writing – review and editing. **CS:** Conceptualization; funding acquisition; methodology; supervision; writing – original draft; writing – review and editing. **PL:** Conceptualization; funding acquisition; methodology; writing – review and editing.

## Acknowledgements

We would like to express our gratitude to the Department of Women’s Health, Women’s University Hospital, and Tübingen University Hospital for their invaluable support and provision of fresh tumor tissue biopsies and corresponding FFPE material. We extend our gratitude to all patients for giving their informed consent for secondary use of residual tissue.

