## Supplementary Data for "Ovarian cancer-on-chip for patient-specific profiling of treatment responses to sequential chemotherapy and PD-L1 blockade"

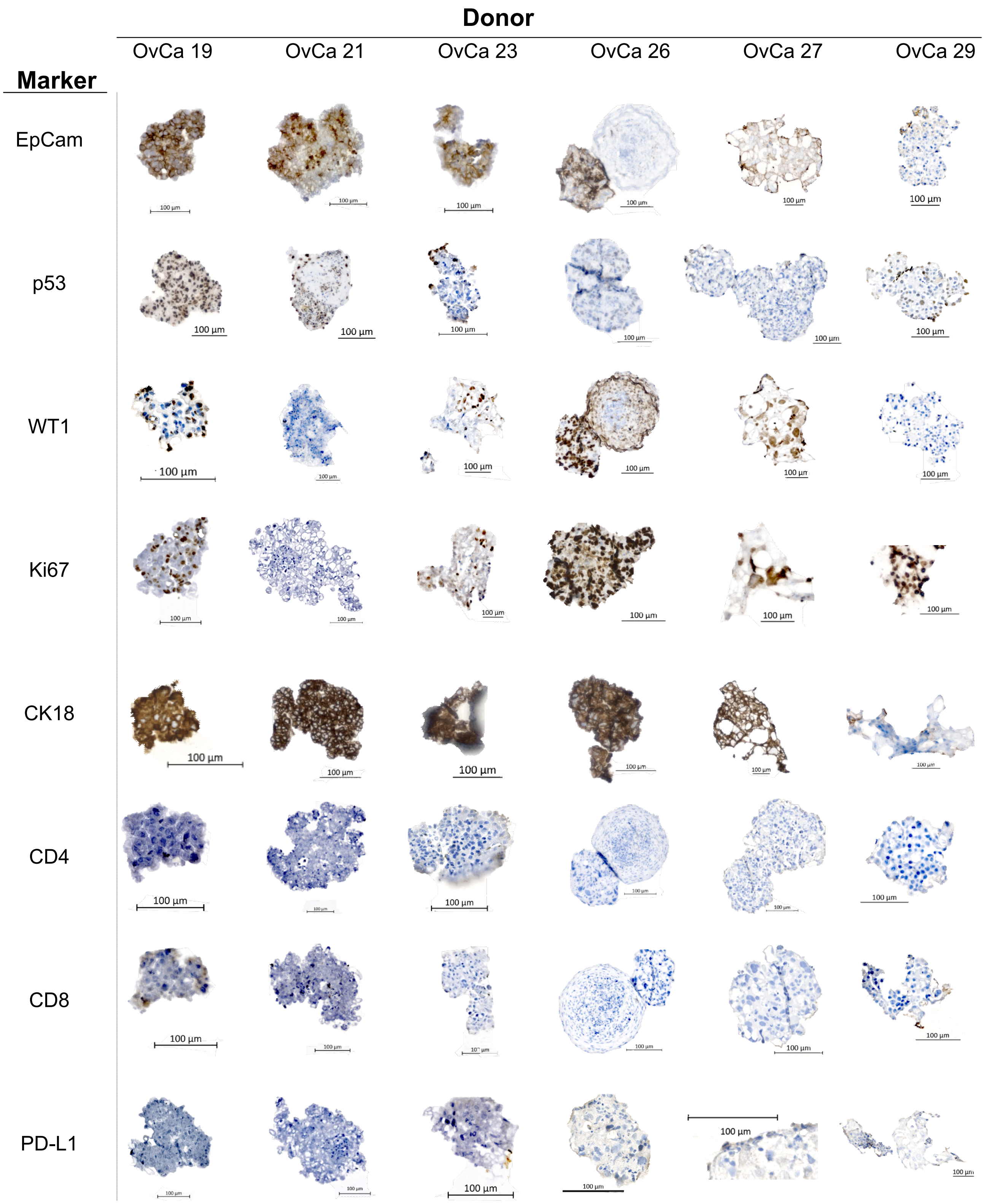


**Figure S1.** Immunohistochemical characterization of OvCa PDM. Representative images of EpCam, p53, WT1, Ki67, CK18, CD4, CD8 and PD-L1 immunohistochemical stainings of different OvCa PDM donors are shown. Positive staining is indicated by a brown color. Nuclei were counterstained with hematoxylin as described in the Methods section and are shown in blue. Scale bars correspond to 100 µm.

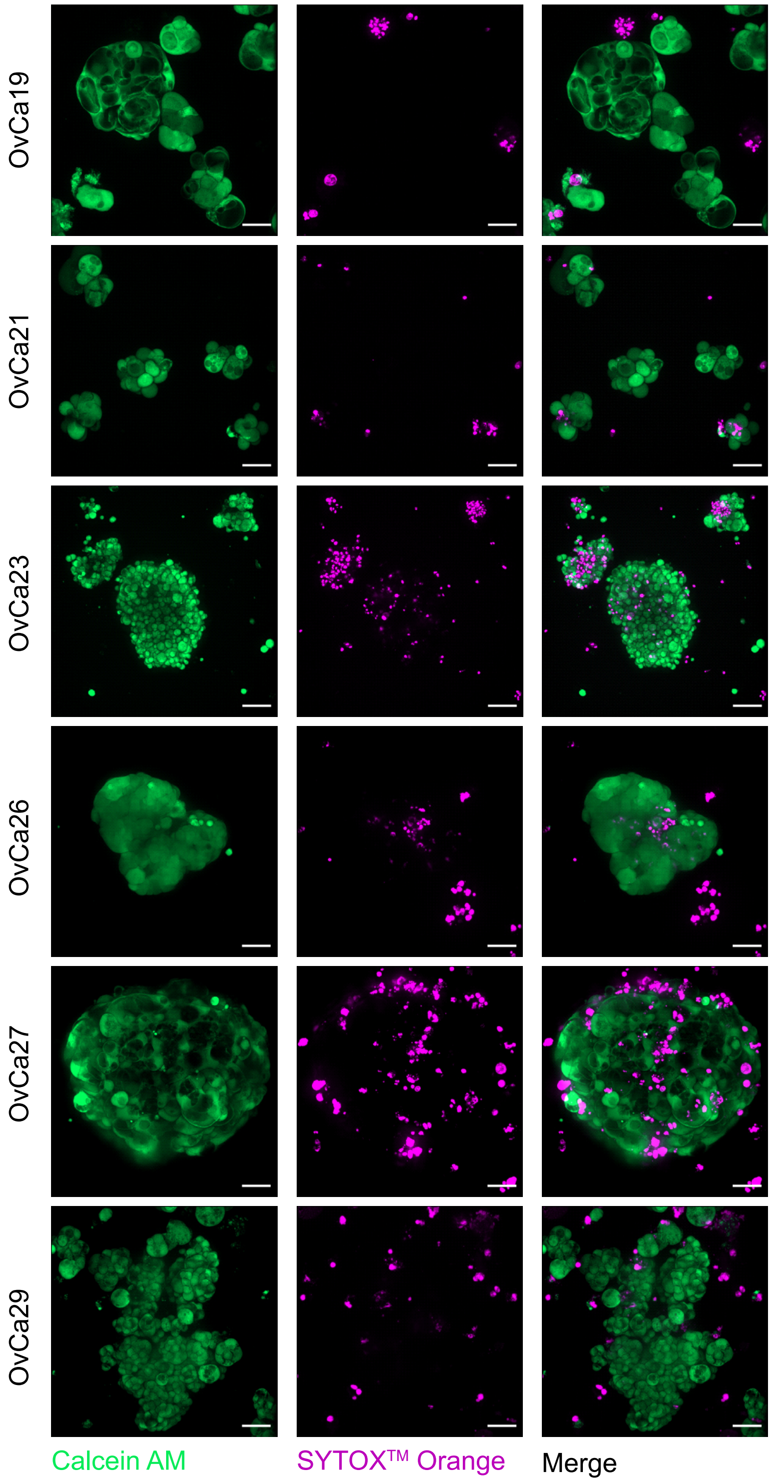


**Figure S2**. Viability staining of OvCa PDM after isolation. Live/dead cell staining was performed for n = 6 PDM donors after PDM isolation. Representative images of live/dead staining are shown for indicated OvCa PDM donors. Live cells are shown in green (stained with Calcein); dead cells are shown in purple (stained with Sytox Orange). Scale bars correspond to 50 µm.

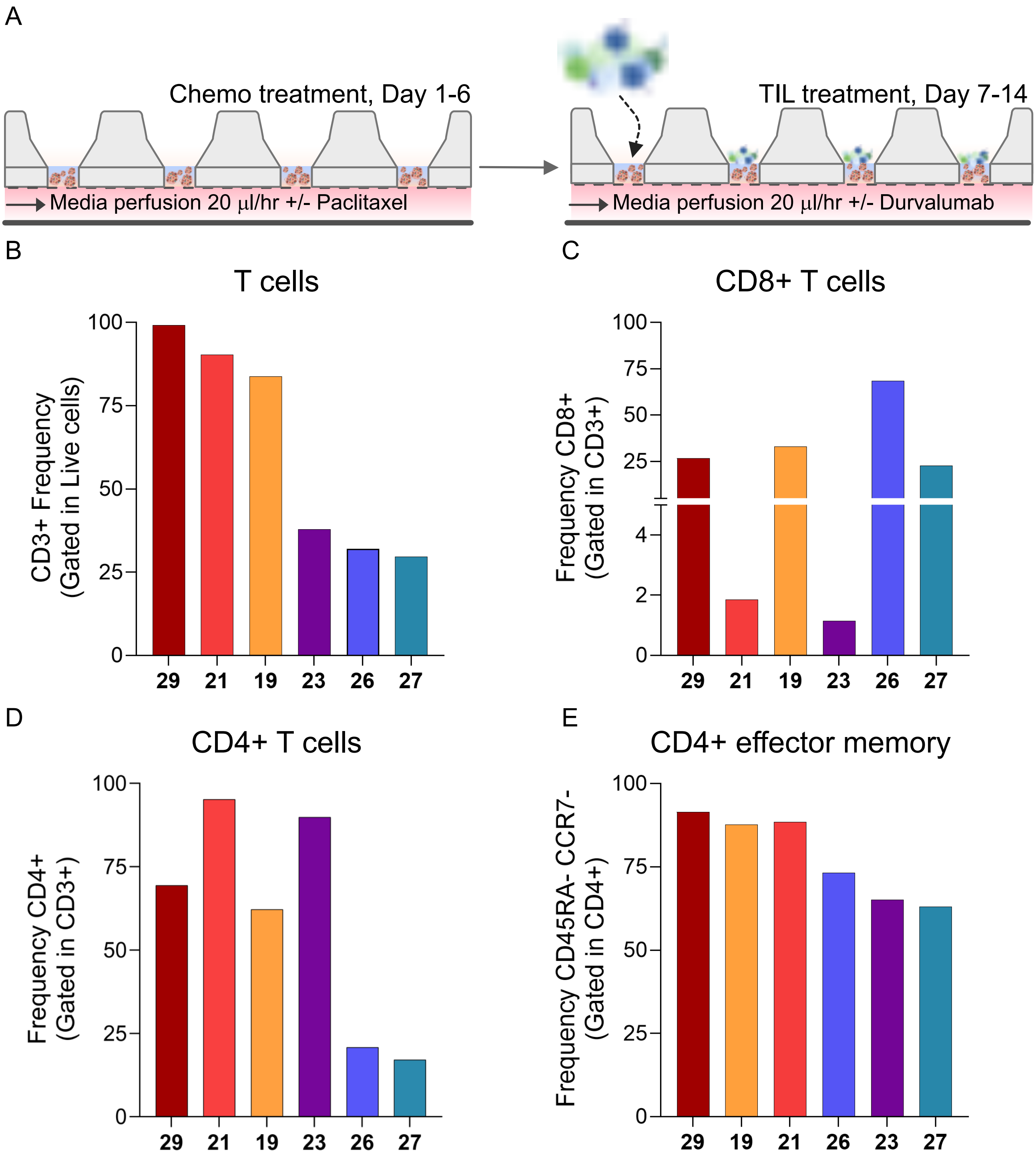


**Figure S3. FACS based phenotypic characterization of autologous ovarian cancer TIL populations**. (**A**) Schematic overview of the experimental procedure. OvCa PDM were seeded in a dextran-based hydrogel. From Day 1 to day 6 PDM were perfused with or without 25 µg/ml paclitaxel. On day 7, autologous TILs (E:T 10:1) were added to PDM and perfused with or without durvalumab until day 14. Effluent was collected every 24h. 50 PDM per chip were seeded, three chips per condition were cultured. Representative flow cytometric characterization of TIL subpopulations used for co-culture experiments, including frequences of (**B**) $CD3^+$ T cells, (**C**) $CD8^+$ T cells, (**D**) $CD4^+$ T cells, and (**E**) $CD4^+$ effector memory T cells.

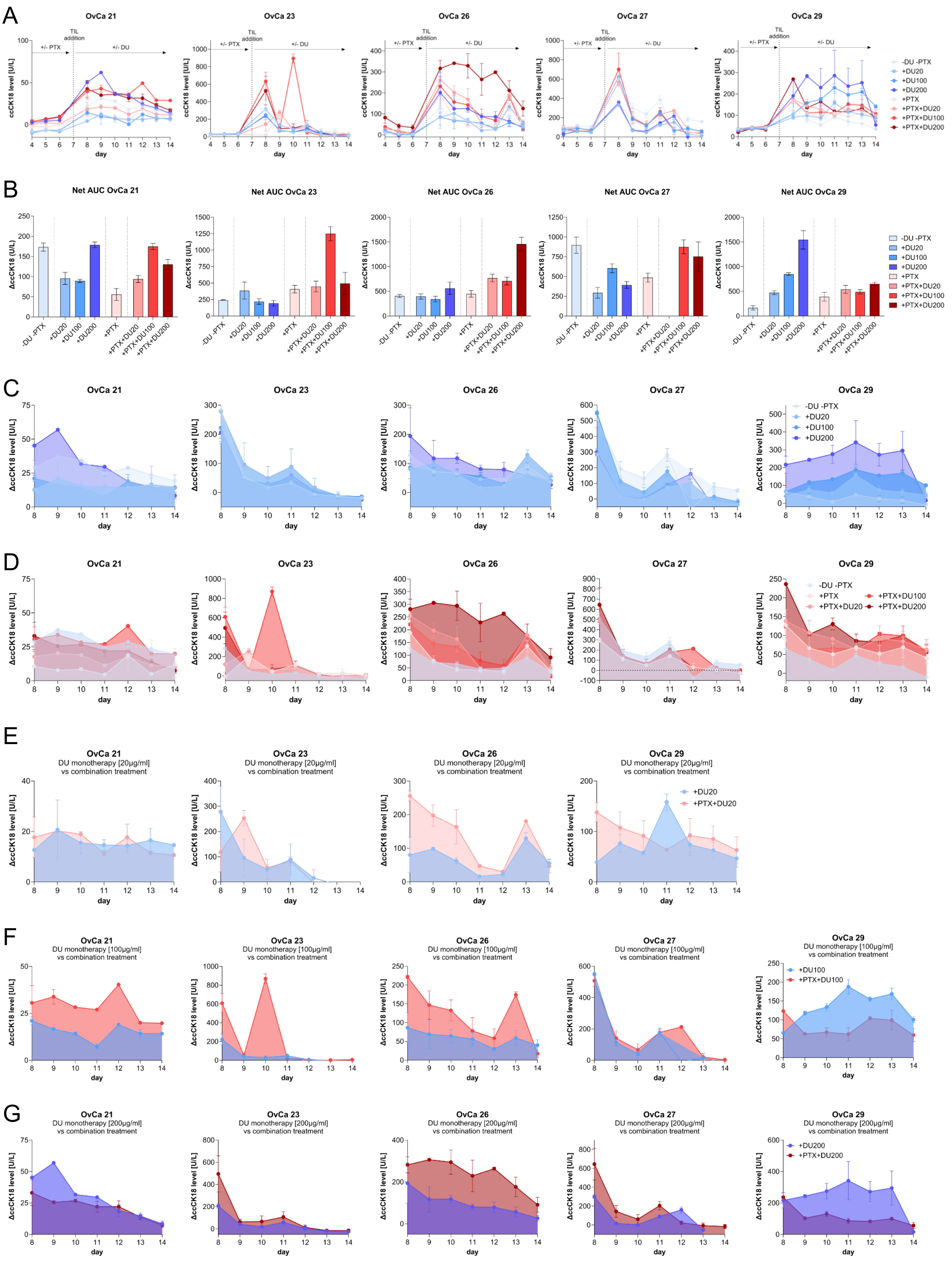

**Figure S4. Functional response profiling of sequential chemo-immunotherapy in patient-derived OvCa-on-chip cultures**. (**A**) Longitudinal ccCK18 kinetics of OvCa21, OvCa23, OvCa26, OvCa27 and OvCa29 PDM-on-chip cultures from day 4 to day 14 across all treatment conditions. (**B**) Treatment-associated tumor cell death following TIL treatment. Values shown represent area under curve (NetAUC) of ccCK18 values, corrected for the baseline (day 6), from day 8 to day 14. (**C–G**) Kinetics of tumor cell death after TIL addition on day 7, as well as additional treatment effects of chemotherapy pretreatment with or without PD-L1 blockade (day 8-14, values corrected for baseline day 6). (**C**) Treatment response to anti-PD-L1 therapy compared to TIL-treatment. (**D**) Treatment response to sequential chemo-immunotherapy (**E–H**) The impact of PTX pretreatment on the efficacy of PD-L1 blockade. Direct comparison of DU treatment with the corresponding combination PTX+DU treatment at DU20, DU100 and DU200, respectively.

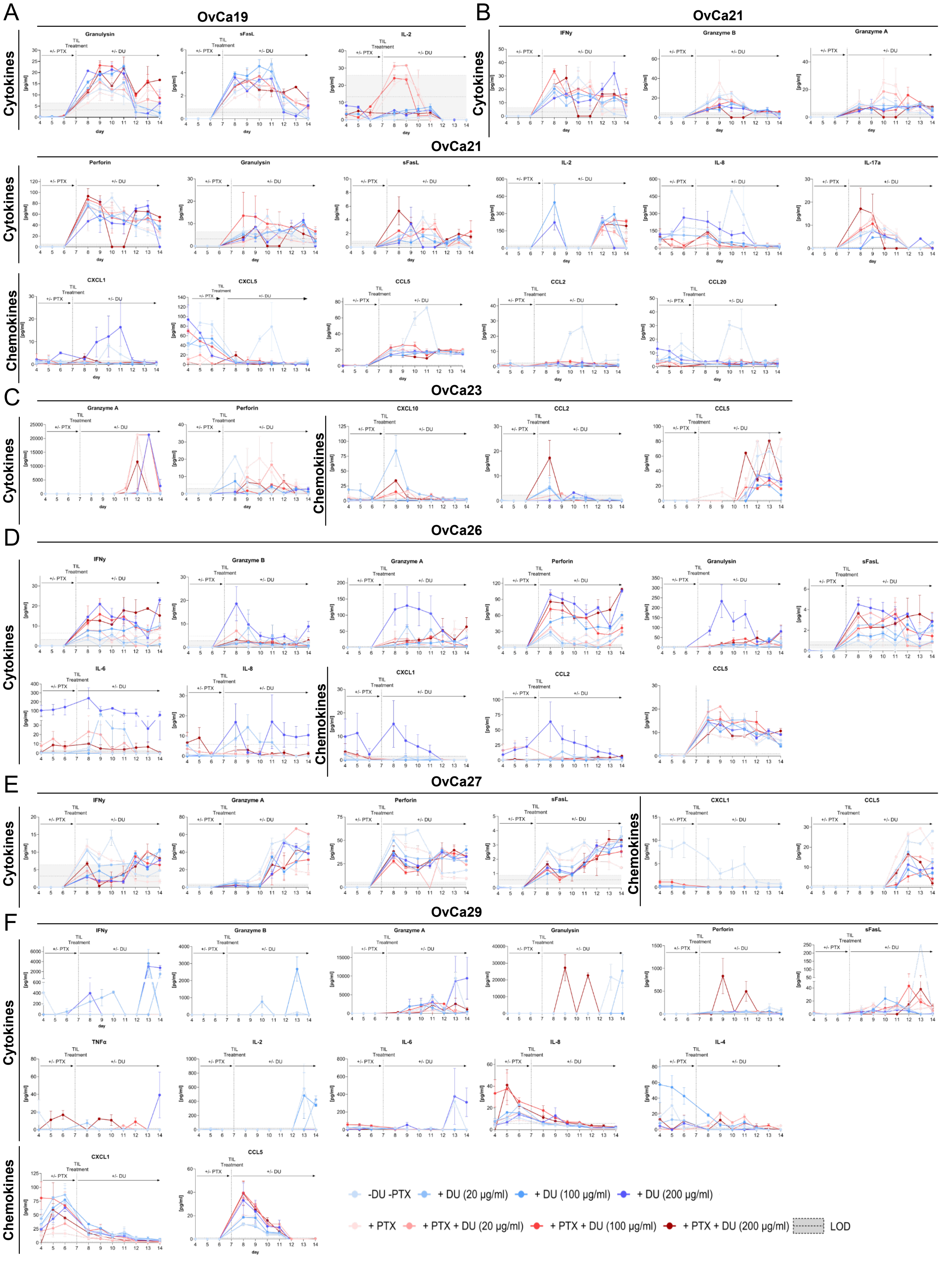


**Figure S5. Longitudinal secretome profiling reveals donor- and treatment-dependent patterns.** Kinetics of cytotoxic TIL effector mediators and chemokines in treated on-chip cultures of (**A**) OvCa19, (**B**) OvCa21, (**C**) OvCa23, (**D**) OvCa26, and (**E**) OvCa29. Concentrations were quantified of chip effluents collected from day 4 to day 14.

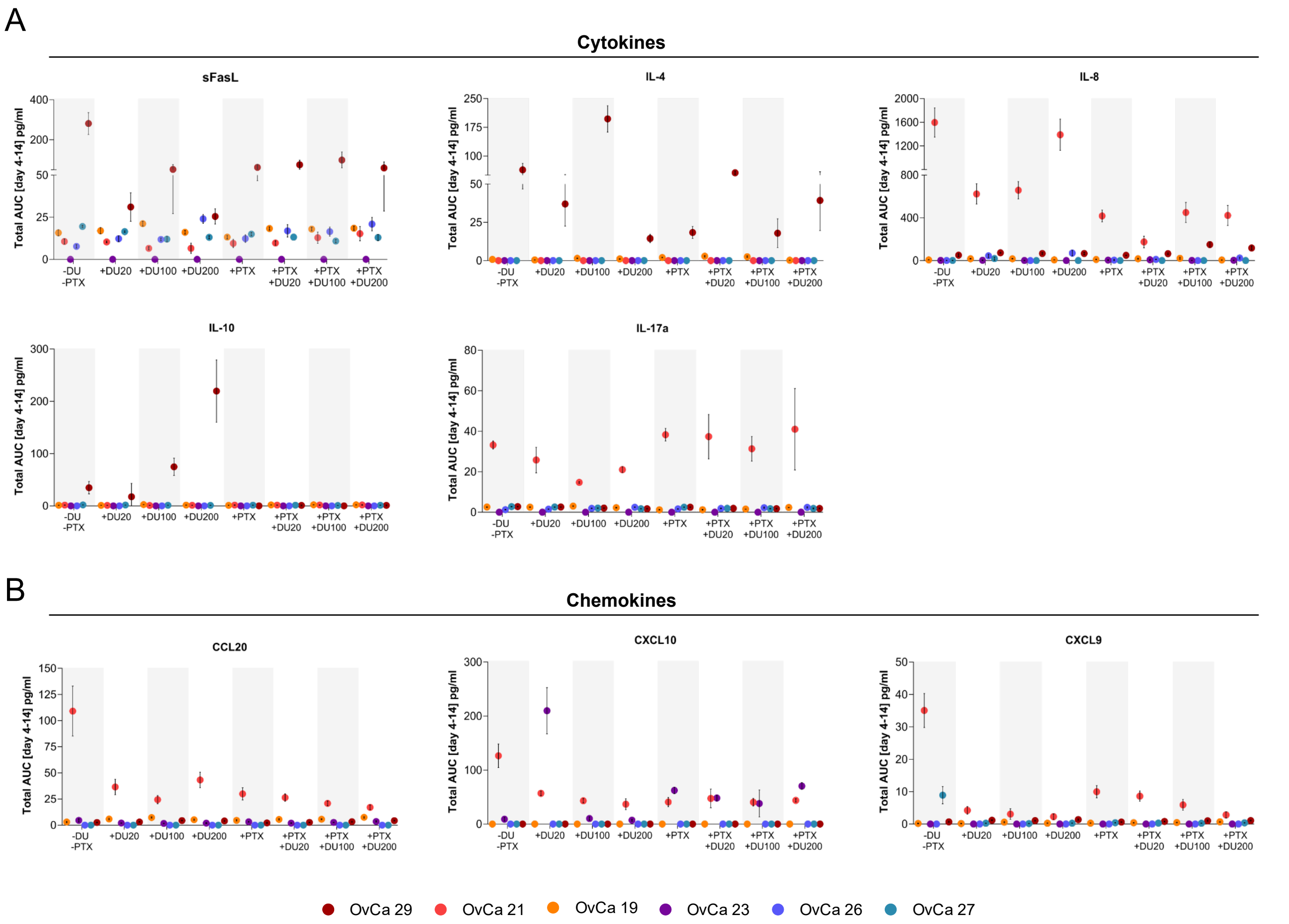


**Figure S6. Cumulative secretome profiles across patient-derived OvCa models and treatment conditions.** (**A**) Cumulative cytokine and (**B**) chemokine profiles shown as Total AUC of cytokine/chemokine concentrations measured from day 4 to day 14. Each color represents one donor.

| | Diagnosis date | OP date | Death date | Last status date | Censoring status | PTX-containing clinical therapy | Clinical therapy context | OS (days) | OS (months) | First dated progression/re currence | Event source | Time to event from diagnosis (days) | Time to event from diagnosis (months) | Time to event from OP (days) | Time to event from OP (months) | First-line response | Maintenance treatment(s) | Metastasis at diagnosis | Clinical course |
|---|---|---|---|---|---|---|---|---|---|---|---|---|---|---|---|---|---|---|---|
| **OvCa 19/21** | 08.11.2021 | 23.11.2021 | 01.03.2022 | 01.03.2022 | Deceased † | No | No paclitaxel-containing regimen documented in the provided table; progress noted. | 113 | 3.7 | | First-line response: Progression (date not specified) | | | | | Progress | None | Yes | Rapid progression/aggressive course |
| **OvCa 21/21** | 09.12.2021 | 09.12.2021 | | 21.03.2023 | censored | No | Radiotherapy documented; no paclitaxel-containing systemic therapy documented. | 467 | 15.3 | | No dated progression/recurrence | | | | | Not documented | None | No documented metastasis | Limited follow-up/no documented progression |
| **OvCa 23/22** | 08.04.2022 | 21.04.2022 | | 23.06.2023 | censored | Yes | Carboplatin/Paclitaxel/Bevacizu mab followed by Bevacizumab + Olaparib maintenance. | 441 | 14.5 | 21.02.2023 | Maintenance response: pulmonary metastasis | 319.0 | 10.5 | 306.0 | 10.1 | Stable; no evidence of metastases | Bevacizumab + Olaparib | Yes | Documented progression after initial disease control |
| **OvCa 26/22** | 30.06.2022 | 02.08.2022 | 18.11.2025 | 18.11.2025 | Deceased † | Yes | Carboplatin/Paclitaxel followed by Niraparib maintenance. | 1237 | 40.6 | | No dated progression/recurrence in provided table | | | | | No evidence of metastases | Niraparib | Yes | No documented progression in provided table; deceased |
| **OvCa 27/22** | 22.06.2022 | 03.08.2022 | 29.05.2025 | 29.05.2025 | Deceased † | No | Carboplatin first-line; later Carboplatin and palliative Doxorubicin documented. | 1072 | 35.2 | 13.03.2024 | Maintenance response: progression LK/HEP/peritoneum | 630.0 | 20.7 | 588.0 | 19.3 | Stable; no evidence of metastases | Niraparib; later carboplatin; later doxorubicin | No documented metastasis at diagnosis | Documented progression after initial disease control |
| **OvCa 29/22** | 10.08.2022 | 17.08.2022 | 24.01.2023 | 24.01.2023 | Deceased † | No | No paclitaxel-containing regimen documented; early progression documented. | 167 | 5.5 | 25.10.2022 | First-line response/date first recurrence: progression peritoneum | 76.0 | 2.5 | 69.0 | 2.3 | Progress peritoneum | None | Yes | Rapid progression/aggressive course |

**Table S1.** Clinical patient data for PDM used in the study

| Antibody | Host Species | Supplier | Dilution |
|---|---|---|---|
| CD3 | rabbit | Agilent Dako (IR503) | RTU |
| CD68 | mouse | Agilent Dako (IR609) | RTU |
| Collagen I | rabbit | Biomol (R31258) | 1:1000 |
| CK18 | mouse | Agilent Dako (IR618) | RTU |
| EpCam | mouse | Agilent Dako (IR63761) | RTU |
| FAPα | rabbit | Biorad (AHP1322) | 1:50 |
| Ki-67 | mouse | Agilent Dako (IR626) | RTU |
| PD-L1 | mouse | Agilent Dako | 1:50 |
| TP53 | mouse | Agilent Dako (IR616) | RTU |
| WT1 | mouse | Agilent Dako (IR055) | RTU |

RTU: Ready-to-use

**Table S2.** Immunohistochemistry primary antibodies, their source and the working concentration

| Antibody | Species | Supplier | Staining | Dilution |
|---|---|---|---|---|
| anti-human PD-1, BV421 | mouse | BioLegend (329920) | extracellular | 1:20 |
| anti-human CD4, BV570 | mouse | BioLegend (300534) | extracellular | 1:20 |
| anti-human CD69, BV650 | mouse | BioLegend (310934) | extracellular | 1:20 |
| anti-human CD39, BV785 | mouse | BioLegend (328240) | extracellular | 1:20 |
| anti-human CD8, PerCP-Cy5.5 | mouse | BioLegend (344710) | extracellular | 1:20 |
| anti-human CD3, FITC | mouse | BioLegend (344804) | extracellular | 1:20 |
| anti-human Tcf1, PE | mouse | BioLegend (655208) | intracellular | 1:20 |
| anti-human CD103, PE-Cy7 | mouse | BioLegend (350212) | extracellular | 1:20 |
| anti-human FOXP3, AF647 | mouse | BioLegend (320114) | intracellular | 1:20 |
| anti-human CD25, AF700 | mouse | BioLegend (302919) | extracellular | 1:20 |
| Zombie NIR™, APC,Cy7 | | BioLegend (423106) | | 1:1000 |

**Table S3.** FACS antibodies, source, manufacturer and working concentrations

| OvCa-on-chip | ccCK18 response pattern | PTX rank | Best DU rank | Best PTX+DU rank | FC PTX vs. matched TIL AUC) | FC DU vs. matched TIL AUC | FC PTX+DU vs. matched DU AUC | TIL-only NetAUC | PTX NetAUC | Best DU condition NetAUC | Best PTX+DU condition NetAUC | matched DU NetAUC (for PTX-DU effect) |
|---|---|---|---|---|---|---|---|---|---|---|---|---|
| **19** | TIL-driven killing;<br>No DU-sensitivity;<br>PTX pretreatment increases DU response (PTX driven) | **+/-** | **+/−<br>(DU200)** | **+++<br>(PTX+DU100)** | 1.1 | 0.7 | 3.7 | 276.3 | 304.9 | 194.4 | 341.6 | -91.11 |
| **21** | Limited/low response; | **+/−** | **+/−<br>(DU200)** | **+++<br>(PTX+DU100)** | 0.3 | 1.0 | 2.0 | 173.2 | 55.8 | 178.5 | 174.7 | 89.17 |
| **23** | No broad sensitivity to DU;<br>PTX-sensitive;<br>Benefit from PTX pretreatment (strongest combination-enhancement profile) | **++** | **++<br>(DU20)** | **+++<br>(PTX+DU100)** | 1.7 | 1.6 | 5.7 | 241.8 | 407.8 | 385.4 | 1250 | 220.1 |
| **26** | DU200 sensitive;<br>No PTX-sensitivity;<br>Benefit from PTX pretreatment (sustained PTX+DU-associated response) | **+/-** | **++<br>(DU200)** | **+++<br>(PTX+DU200)** | 1.1 | 1.4 | 2.6 | 406.1 | 447.5 | 560.6 | 1456 | 560.6 |
| **27** | TIL-driven killing;<br>No PTX or DU enhancement | **+/−** | **+/−<br>(DU100)** | **++<br>(PTX+DU100)** | 0.5 | 0.7 | 1.4 | 896.4 | 485.8 | 604 | 875.7 | 604 |
| **29** | PTX-sensitivity;<br>Strong DU-sensitivity (doese-dependent)<br>PTX attenuates DU-associated response | **+++** | **+++<br>(DU200)** | **+/-<br>(PTX+DU200)** | 2.4 | 9.5 | 0.4 | 163 | 390.3 | 1544 | 649 | 1544 |

FC: Fold change
FC ≥ 2 (+++) = clear treatment-associated enhancement
FC ≥ 1.25 (++) = modest enhancement
FC < 1.25 (+/-) = no relevant enhancement

**Table S4. Ranking of treatment responses in OvCa-on-chip platforms based on ccCK18.** Tumor cell death was assessed by NetAUC of ccCK18 values day 8-14). Treatment responses were assigned based on calculated fold change (FC) values. FC was calculated using the NetAUC of the following matched treatments: PTX vs. TIL (PTX response); DU vs. TIL (DU response); and PTX + DU vs. DU (PTX pretreatment effect on DU). The relevant NetAUC values used in the calculation are listed.
